# Single cell Correlation Analysis (SCA): Identifying self-renewing subpopulation of human acute myeloid leukemia stem cells using single cell RNA sequencing analysis

**DOI:** 10.64898/2026.02.27.708585

**Authors:** Yoonkyu Lee, Wen Wang, Timothy K Starr, Klara E. Noble-Orcutt, Chad L. Myers, Zohar Sachs

**Author notes:** Corresponding authors: Zohar Sachs, 420 Delaware St SE, MMC 806, Minneapolis, MN 55455.

## Abstract

Leukemia stem cells (LSCs), a rare and self-renewing subpopulation, drive Acute myeloid leukemia (AML) relapse and therapy resistance. While single-cell gene expression profiling (scGEP) offers high-resolution insights into LSC biology, existing computational tools rely on relative and arbitrary similarity metrics. This limits their ability to definitively identify true biological cell identities or compare rare cell populations across independent datasets. To overcome these limitations, we developed Single cell Correlation Analysis (SCA), a novel computational method that utilizes a permutation-based false discovery rate (FDR) framework and common background datasets to establish standardized, statistically rigorous thresholds of similarity. We applied SCA to query human AML scRNA-seq datasets using an experimentally validated murine scGEP of LSC self-renewal as our reference. SCA demonstrated superior specificity and precision, maintaining low false positive rates compared to existing reference-based annotation tools. Using SCA, we successfully identified cells expressing a conserved self-renewal program (scaLSC-SR) in both adult and pediatric human AML samples. These scaLSC-SR cells share classical LSC immunophenotypic markers (e.g., CD34, CD96, CD200) and exhibit transcriptomic hallmarks of LSC biology, including evasion of apoptosis and immune responses. Furthermore, this self-renewal program is significantly enriched in poor-risk genetic subtypes, specifically *TP53* and *NRAS* mutated AML. Finally, we derived a 28-gene signature (LSC-SR28) from human AML patients that accurately captures LSC stemness and is highly prognostic across multiple independent clinical cohorts. SCA provides a robust, statistically principled framework for identifying rare and biologically meaningful cell populations across heterogeneous single-cell datasets. By successfully identifying a functional self-renewal profile to human AML, SCA reveals conserved, clinically relevant LSC populations that drive relapse and therapy resistance across different age groups.

## Introduction

Acute myeloid leukemia (AML) is an aggressive blood cancer with a two-year survival rate of less than 50%(1). Although standard chemotherapy induces complete remission in most patients, more than 50% relapse and only 20–30% survive beyond two years(2).

Leukemia stem cells (LSCs) are defined as the cells able to recapitulate leukemia, a property known as self-renewal capacity. LSCs are a rare subpopulation of AML cells that drive therapy resistance and relapse(3, 4). Conventional chemotherapy targets rapidly dividing cells, but LSCs are quiescent and evade treatment(5). Therefore, understanding the biology and molecular mechanism of LSC self-renewal is a critical step to eliminate AML relapse and reduce disease mortality.

LSCs are often defined based on their cell surface protein expression patterns, termed their immunophenotype. However, immunophenotypic markers are poor surrogates for LSC function. Defining the molecular basis of self-renewal would provide a more functionally-relevant representation of LSCs. Therefore, several groups have sought to define the transcriptional profile of LSCs(6–8). In these studies, human AML samples sorted according to immunophenotypic LSC markers were transcriptionally profiled to define the gene expression profile (GEP) of each immunophenotypic compartment. In some of these studies, the stem cell compartment was functionally assayed using the gold-standard *in vivo* murine reconstitution assays(6, 8, 9). These studies were instrumental at establishing the transcriptional profile of self-renewal in AML and demonstrated that the LSC stemness gene expression signature is highly prognostic; strongly correlating with poor patient outcomes (8). However, the sorted leukemia fractions profiled in these studies were not pure LSC fractions. These fractions were enriched for LSCs but also contained cells that lack self-renewal capacity.

Single-cell transcriptional studies have also been used to interrogate the molecular biology of self-renewal in LSCs. Previous study used a machine learning approach and single cell genotype and gene expression profile to distinguish malignant from normal cells based on their scRNA-seq features (10). This analysis defined LSCs based on their transcriptional profiles and showed that these transcriptionally-defined LSCs exhibited dysregulated transcriptional programs compared to normal HSCs, and their GEPs were associated with poor prognosis. However, the LSCs in this study were defined based on their GEPs and were not experimentally validated. Another study utilized an lentiviral microRNA-126 (*miR-126*) reporter to prospectively identify and track functionally active LSCs in *NPM1*-mutated human AML patient-derived xenograft models before and after in vivo chemotherapy (11). The authors performed bulk gene expression profiling on flow-sorted miR-126-high and miR-126-low populations, deriving a functional LSC gene signature. Therefore, the functional LSC gene expression profile was not directly derived at the single-cell level. Instead, the authors mapped this bulk-derived signature onto scRNAseq to locate these cells within the broader leukemia population.

In our prior work(12), we sought to define the scGEP of pure LSCs that demonstrate self-renewal capacity *in vivo*. We studied the LSC-enriched compartment of the *Mll-AF9/NRASG12V* model of murine AML that we had previously defined as (Mac1^Low^cKit^High^,Sca-1^High^ (MKS)(13). scRNA-seq identified three distinct GEPs within the sorted MKS compartment. These GEPs were delineated by genes encoding the cell surface markers, CD36 and CD69. To determine whether these distinct GEPs reflect functional heterogeneity within the MKS subset, we analyzed the proliferative and self-renewal capacity of MKS cells according to their CD36 and CD69 protein expression.

The self-renewal capacity of sorted MKS subsets was evaluated using *in vitro* colony-forming assays and *in vivo* reconstitution assays. The CD36^Low^CD69^High^ MKS subgroup showed the highest self-renewal capacity, while CD36^High^CD69^Low^ MKS cells failed to self-renew in all assays. Additionally, *in vivo* proliferation tracing revealed that the CD36^High^CD69^Low^ MKS subgroup was highly proliferative. In contrast, the self-renewing CD36^Low^CD69^High^ subgroup was poorly proliferative. These data demonstrate that the scGEPs reflect critical functional subsets within the LSC-enriched compartment of murine AML. Furthermore, these single-cell transcriptional analyses demonstrate that self-renewal and rapid proliferation are mutually exclusive within the LSC compartment, as was previously demonstrated in normal HSCs(14).

Our murine studies defined the leukemia self-renewal GEP at the single-cell level and demonstrated that scGEPs reveal important functional leukemia subsets that are imperceptible using bulk methods. Here, we seek to determine whether human AML samples harbor cells that express this scGEP of self-renewal. Defining these cells at the single-cell level in would permit a more precise understanding of the molecular basis for self-renewal and define potential therapy targets in human AML. However, existing scRNA-seq analysis methods cannot definitively resolve whether a GEP of interest is expressed in a query dataset.

Numerous computational tools have been developed to analyze scRNA-seq data, but existing methods are limited by relative similarity metrics. While they can rank which cells are most similar to a reference, they cannot definitively assess true biological cell identity or confirm if a gene expression profile is truly recapitulated in a query dataset. A major goal among existing tools is to annotate a query dataset using an established reference dataset. In this process, cell types in the query dataset are assigned based on their similarity to reference-defined cell types (15–19). These reference-based methods typically rely on correlation coefficients(16, 17, 20, 21) or algorithm-specific scores(15, 18–21) for measuring GEP similarity. Cell type annotations are then assigned using either arbitrary similarity thresholds (e.g., Pearson’s correlation coefficient > 0.7) or relative criteria (selecting most similar cells or highest scoring cells when multiple reference cell types were used). However, these approaches present several limitations, including unknown false positive rates and little justification for threshold values. Therefore, these methods are unable to discern what level of similarity reflects true biological or functional similarity.

To address these limitations, we developed Single cell Correlation Analysis (SCA). SCA is a scRNA-seq analysis method that uses statistical significance to define a threshold of similarity and providing a quantifiable assessment of biological and functional similarity. To establish a statistically-defined threshold of similarity, SCA generates a null distribution by randomizing the dataset. SCA then incorporates a permutation-based false discovery rate (FDR) framework to statistically assess the significance of the similarity of a query cell to a reference GEP (relative to the null distribution). In addition, SCA is designed to directly compare query cells from multiple and unrelated studies. For this case, SCA uses a common background dataset to generate a null distribution, which is then applied to each query dataset to establish a universal threshold across studies. This step enables consistent and comparable results across datasets. This robust statistical framework makes SCA particularly suited for identifying rare, biologically significant populations such as self-renewing LSCs with high statistical confidence.

We compared SCA to other scRNA-seq analysis approaches using published datasets of experimentally validated scRNA-seq data of normal human and murine HSPCs. These comparisons reveal that SCA’s statistically-defined FDR optimizes for high specificity and minimizes false positive rates, while maintaining high sensitivity. Next, we applied SCA to published scGEPs of human AML samples and found that these samples express the scGEP of self-renewal that we defined and experimentally validated in our murine model (scaLSC-SR cells). We compared the scGEPs human scaLSC-SR cells from adult and pediatric patients and characterized their expression of functionally annotated gene expression profiles and cell surface marker genes. Finally, we showed that the GEPs of scaLSC-SR are preferentially expressed in poor risk AML subtypes and the expression of this GEP is associated with poor survival in three independent transcriptional datasets of human AML.

## Methods

### Data Preprocessing

All analyses were performed using R (v4.4.1) software. scRNA-seq data were acquired for human normal HSPC datasets (10, 22), murine normal HSC(23), murine LSC(12), adult AML(10) and pediatric AML(24). scRNA-seq data with raw read count or unique molecular identifier (UMI) count data were preprocessed using scran (v1.32.0) package in R. Cells with more than 500 detected genes and less than 15% mitochondrial gene content were retained for downstream analysis, and genes expressed in at least 10 cells were included in further analyses. Each cell was normalized based on library size factors and log-transformed with a pseudocount of 1. For visualization of single-sample scRNA-seq datasets, PCA was performed using the top 15% of highly variable genes and normalized data, followed by tSNE on the PCA embeddings. The runTSNE function in the scater (v1.32.1) R package was used for the analysis. Similarly, scRNA-seq data from multiple samples(10, 24), were batch-corrected using the fastMNN function in the batchelor (v1.20.0) R package with log-transformed data and the top 5000 highly variable genes. Following batch correction, PCA was performed on the corrected expression values, and then tSNE was subsequently performed using the PCA embeddings.

The raw bulk RNAseq read count data were obtained from the Beat AML(25), TCGA LAML(26, 27), and sorted AML fraction(8, 9) data. Genes that were expressed in more than 5 samples were kept for the downstream analysis.

The raw microarray CEL data were obtained from the Gene Expression Omnibus (GEO; GSE6891 and GSE37642). The data were processed and normalized to generate probe-level expression matrices using ReadAffy function from affy R package (v1.82.0). GSE6891 was captured in GPL570 platform and GSE37642 was captured in both GPL570 and GPL96 platforms. Both platforms were combined in GSE37642 using the common probe sets. A standard background correction and normalization process were performed with rma function in affy R package. Batch correction was performed on combined GSE37642 using Combat function in sva (v3.52.0) R package. Probe matrix is then converted to gene-level expression matrices using provided probe to gene map in the data.

### Inclusion and exclusion criteria in bulk RNA Seq dataset analysis

We analyzed two publicly available independent AML data sets, Beat AML (25) and TCGA LAML(26, 27) (including Beat AML data updates(25)).

<u>Beat AML</u>. We included every sample that has both whole exome sequencing (WES) and RNAseq data available (n=615). <u>TCGA LAML</u>. We included every sample that has both DNA sequencing and RNAseq data available (n=163). DNA sequencing was available as either WES or whole genome sequencing (WGS). The LAML samples in the TCGA are all diagnostic.

### Genetic Data of bulk RNAseq datasets

#### Beat AML

Mutation assessments were reported by the Beat AML dataset and data updates(25, 28) based on WES. WES coverage data is not available for this dataset. *TP53* mutations are reported at variant allele frequency (VAF) as low as 0.5% in both datasets indicating sensitivity in this range. Karyotype and fluorescence *in situ* hybridization (FISH) data is reported for this dataset as well. CEBPA, FlT3-ITD, NPM1, RUNX1, RUNX1-RUNX1T1, ASXL1, TP53 mutation annotations were taken from the clinical data previous study provided.

#### TCGA LAML

Mutation assessments were reported by the TCGA LAML dataset (26) either WES (which produced an average of 30.54x coverage) or WGS (which produced an average of 167.50X coverage).

### Single cell Correlation Analysis (SCA)

SCA identifies individual cells in query dataset that expresses a specified reference gene expression profile. To assess the statistical significance of the similarity between each query cell and the reference profile, SCA employs permutation tests to derive empirical p-values and estimates the false discovery rate (FDR) to correct for multiple hypothesis testing. SCA can utilize any reference profile, whether derived from bulk RNAseq or scRNA-seq. FDR thresholding provides a systematic way to control proportion of false discoveries, maximizing significant findings while minimizing false positive rates. Additionally, SCA offers adaptive thresholding for cell type assignments, applying a more stringent FDR when the evidence is weak and a more permissive FDR when the signal is strong. This process defines an absolute threshold for a level of similarity that exceeds what may be seen by random chance.

While SCA can use any provided reference profile, SCA provides a method to construct a reference profile from scRNA-seq data through differentially expressed gene (DEG) analysis, using scoreMarkers function from the scran (v1.32.0) R package. Based on the DEG result, SCA ranks genes by AUC and takes the top n/2 and bottom n/2 genes by Cohen’s d log-fold change values to construct the reference profile, where n is the user-defined number of genes to include. This pipeline for constructing reference profile was used in all scRNA-seq analyses. SCA assigns each cell a similarity score by calculating Spearman’s correlation coefficient (SCC), observed SCC, between the reference profile vector and each cell in the query single cell gene expression data. SCA then requires a background single cell gene expression dataset to generate randomized data, which is used to compute a null distribution for SCC statistics 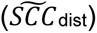 by comparing the reference profile with pseudo-cells from the randomized data. The randomized data is generated by randomly permuting expression values of each gene across cells in the background data. Null SCCs are calculated between the reference profile vector and each pseudo-cell in the randomized data. This process was repeated 1,000 times to generate a robust null SCC distribution in all the analyses in this paper. The background data can be either the query data itself or another scRNA-seq dataset from same tissue type, provided it contains diverse cell types. If no external background data is provided, the query dataset will be used. We evaluated the use of different background datasets for SCA, with detailed methods described in the ‘SCA Input Background Data Evaluation’ section. SCA uses only the genes that are shared across reference profile, query data and background data to generate observed SCCs and null SCCs. With these observed SCCs and the null SCC distribution, SCA calculates empirical p-values using empPvals function from the qvalue (v2.36.0) package in R (29). Finally, false discovery rate (FDR) q-values are estimated from the empirical p-values using qvalue function.

SCA was tested on normal human HSPC scRNA-seq datasets(10, 22) to identify cells that express murine HSC self-renewal (HSC-SR) single cell gene expression profiles. A murine HSC self-renewal scRNA-seq data(23) was used as a reference profile. For each human HSPC data, the query dataset was used as background data.

SCA was applied to 16 adult(10) and 13 pediatric AML scRNA-seq datasets(24) to identify AML cells expressing murine LSC self-renewal (scaLSC-SR) and proliferation (scaLSC-Pro) gene expression profiles at the single-cell level. The murine LSC self-renewal and proliferation gene expression profiles from(12) were used as reference profiles. All 16 adult AML scRNA-seq datasets were used as background data for querying adult AML, and 13 pediatric AML scRNA-seq datasets were used as background data for querying pediatric AML datasets.

To compare adult and pediatric AML, cell type annotations defined from diagnostic adult AML scRNA-seq data were transferred to pediatric AML scRNA-seq datasets. The SingleR (v2.6.0) R package was used for transferring cell type annotation, with default settings except for the use of the Wilcoxon rank sum test as the marker detection mode to identify top markers.

### SCA input background data evaluation

SCA defaults to using query data as background data when no external background is provided. SCA was designed to utilize external background data, making it particularly useful for comparing results across multiple scRNA-seq dataset from different samples or conditions. When SCA uses one external background data to generate a single SCC null distribution 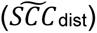, statistical significance of individual cell’s correlation coefficient from different dataset is assessed against the same 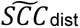. This approach facilitates consistent and effective comparisons across datasets. External background data can include scRNA-seq data from the same or relevant tissue types, ideally encompassing a broader or similar range of cell types. In this analysis, each human HSPC query dataset was comprised of diverse cell types and was therefore heterogeneous enough to serve as its own background dataset. Because SCA employes Spearman’s correlation, a rank-based correlation, SCA is robust to the batch effect. For the same reason, data from different sequencing technologies and quantification methods (e.g., UMI and read counts) may be used as background data. Importantly, SCA generates randomized background data preserving the variance of each gene within the background dataset. However, sorted scRNA-seq data (either computationally or immunophenotypically) that enriches for a specific cell type of interest (reference profile) may not provide adequate 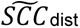.

We evaluated how different background data can affect SCA results, including sorted normal bone marrow (BM) data. First, we assessed the level of cell type heterogeneity required for background data in SCA (**Supplemental Figure S2**). Next, we tested background dataset derived from relevant tissues that differed in sample origin or single cell capture technology and gene expression quantification methods (**Supplemental Figure S5-S7**).

To evaluate the impact of background data cell type heterogeneity on SCA results for identifying LT-HSCs and HSCs in Velten’s HSPC dataset and HSCs in van Galen’s HSPC dataset, we generated and tested various background datasets. For querying Velten data, different background datasets were created by sequentially adding cell populations from the original data: LT-HSCs alone; LT-HSCs and other HSCs; LT-HSCs, other HSCs, and multipotent progenitors (MPPs); LT-HSCs, other HSCs, MPPs, and common myeloid progenitors (CMPs), and all HSPCs. Similarly, for querying van Galen data, we tested three background datasets: HSCs alone; HSCs and progenitors; and all van Galen BM5 HSPC populations. SCA performed 1,000 permutations for each analysis. To assess cell type heterogeneity in the data, we calculated the percentage of variance explained by principal components (PCs) from PCA and used ROGUE(30) statistic, an entropy-based statistic designed to measure cell population purity.

Next, SCA results were evaluated using background datasets from different sample origins, single-cell capture technologies, and gene expression quantification methods. For querying Velten HSPC data, we compared SCA results using the following background dataset: Velten HSPC dataset; van Galen unsorted BM cells; van Galen HSPCs; and van Galen HSCs. Notably, van Galen’s unsorted BM cells originated from a different donor than the HSPCs and HSCs samples, with samples collected from one donor. Similarly, for querying van Galen HSPC data, we tested SCA results using the following background datasets: van Galen’s HSPC dataset itself; van Galen’s HSCs; van Galen’s unsorted BM cells; and Velten’s HSPC dataset. SCA performed 1,000 permutations for each analysis.

### Cell type classification evaluation

Classification performance of SCA was compared with scmap-cluster (v1.26.0), SingleR (v2.6.0), ScType, clustifyr (v1.16.2) and AUCell (v1.26.0). These methods are capable of using a single reference profile vector to identify a cell type of interest and were chosen for consistent comparisons with SCA. DEGs for cell type of interest in scRNA-seq data were generated using the SCA provided method, and the same DEGs were used as the reference profile for all methods tested. The evaluation focused on classifying hematopoietic stem cell (HSC), a rare cell type that self-renews itself and gives rise to differentiated cell types, in two normal human HSPC datasets(10, 22).

Performance was tested with varying number of DEGs. Other methods were fine-tuned with different hyperparameters, and the highest F1 score achieved by each method was reported.

Performance was evaluated in three different ways: intra-data, Inter-data and Inter-species data evaluations.

For intra-data evaluation, a 5-fold cross-validation method was used to assess the performance of each algorithm. The data were randomly divided into five folds; one-fold was used to generate the reference profile (train), while the remaining four (test) were used to test performance. This process was repeated five times, iterating through the train-test folds each time. HSCs in train fold was used to generate reference profile. The same train-test folds were used across all methods, and median value of evaluation metrics were reported.

In the inter-data evaluation, the Velten HSPC data was used as the query dataset because its cell type annotations were based on immunophenotypic markers, which are conventionally used to study human HSPCs, rather than on transcriptomics-based annotation. The HSC reference profile was derived from van Galen data and tested on the Velten HSPC data.

Inter-species data testing was conducted using mouse HSPC scRNA-seq data (23) as the reference and human HSPC datasets as query. A previous study functionally validated mouse HSC engraftment *in vivo* and traced the engrafted quiescent HSCs in the scRNA-seq data using lentiviral cell barcoding. The engrafted quiescent mouse HSC population served as the reference profile, and Velten human HSPC data was used as query data.

The final evaluation assessed each method’s ability to avoid falsely identifying cells outside the cell type of interest, which would result in the including unwanted cells. The false positive rate (FPR) was measured for each method using the Velten HSPC scRNA-seq data, first with the HSC population removed and then with both HSC and multipotent progenitor (MPP) populations removed.

### Performance evaluation metrics

Four different metrics were used to assess the performance of different methods; precision, sensitivity, F1 score and false positive rates (FPR). Precision is the proportion of true positives out of all the positive predictions (true positives/ (true positives + false positives)). Sensitivity is the proportion of true positives out of all actual true positive cases (true positives / (true positives + false negatives)). F1 score is the harmonic mean of precision and sensitivity, and this metric balances the importance of precision and sensitivity. False positive rate is the percentage of false positives out of all negative cases (false positives/ (false positives + true negatives)).

### Gene expression analyses

For differentially expressed gene (DEG) analysis in single sample scRNA-seq data, scoreMarkers function was used from the scran R package (1.34.0). AUC values from the results were used to rank differentially expressed genes. For DEG analysis for multi-sample scRNA-seq datasets, a conventional bulk RNAseq DEG analysis method was applied on pseudo-bulk transformed scRNA-seq AML datasets. First, unique molecular identifier (UMI) counts in the scRNA-seq data were aggregated by cell type within each sample and across samples to create pseudo-bulk UMI count data. This pseudo-bulk data was then used to perform DEG analysis with the edgeR(31) package (v.4.2.1) using glmQLFit and glmQLFTest functions. Gene set enrichment analysis(32)(GSEA) was performed using the clusterProfiler package (v4.12.6) with gene sets curated from the Molecular Signatures Database (MSigDB v2023.2.Hs, https://www.gsea-msigdb.org/gsea/msigdb) and gene sets manually curated from literature of HSC and LSC self-renewal functions(12).

### Gene expression profile deconvolution

Bulk AML gene expression profile datasets were deconvoluted to estimate the abundances of the previously characterized AML cell types(10), including the cells expressing murine LSC self-renewal (LSC-SR) and proliferation (LSC-Pro) gene expression profile identified using SCA. The CIBERSORTx(33) web portal was used for the deconvolution analysis. Count-per-million (CPM) values from 16 adult AML scRNA-seq with 21 cell types (LSC-SR, LSC-Pro, HSC-like, Prog-like GMP-like, ProMono-like, Mono-like, cDC-like, B, CTL, GMP, HSC, Mono, NK, Plasma, ProB, ProMono, Prog, T, cDC, earlyEry, lateEry, pDC) were used to generate a signature matrix. Deconvolution was performed on Beat AML and TCGA data using CIBERTSORTx default settings.

The S-mode batch correction method was used, and Absolute mode was applied. For each sample, CIBERSORTx generated absolute scores for cell type abundances, which were then normalized to sum to 1, so that each cell type abundance score reflects the proportion within the sample. Wilcoxon rank sum tests were performed to compare LSC-SR compositional differences between different mutational groups.

### LASSO Cox regression

The association between LSC-SR gene signatures derived from AML scRNA-seq and overall survival in AML patients was assessed across four independent cohort datasets (25, 26, 34, 35). A similar approach for deriving functional LSC gene signature score from bulk RNAseq data has been described previously(8). Initially, DEG analysis was performed using the pseudo bulk method described above to identify LSC-SR DEGs from adult AML scRNA-seq data. LSC-SR DEGs with a fold change of 2 and FDR<0.05 (313 genes detected across GSE6891(35), GSE37641(34), Beat AML(25), and TCGA LAML(26)) were used to evaluate the association with overall survival (OS). A Cox proportional hazards model with LASSO regularization (LASSO Cox regression) was applied using the glmnet (v4.1-8) R package. Publicly available gene expression profiles from 537 diagnostic patient samples (GSE6891) were used to train the LASSO Cox model. Survival data were provided by the authors Thirty-four cases, including myelodysplastic syndrome subtypes MDS-RAEB/RAEB-t (17 cases), cases lacking OS data (16 cases) and missing data point (1 case), were excluded from the training set. This resulted in 503 cases being included for analysis. The LASSO Cox model was trained using 10-fold cross-validation method, and the hyperparameter λ was chosen to maximize the Harrel’s C-index, a measure of how well the model predicts the order of survival time. In the final model, 28 genes were selected, and their weighted gene expression were combined to assign an LSC-SR28 signature score to each patient sample. The LSC-SR28 model was tested on three individual cohort data - GSE37641, Beat AML and TCGA LAML. Each cohort was stratified into high and low LSC-SR26 score groups based on the optimal score threshold, which was determined to maximize log-rank statistics for survival outcomes using the surv_cutpoint function from the survminer (v0.4.9) R package. Kaplan-Meier survival analysis was then performed to compare survival outcomes between high and low LSC-SR28 score groups, with log-rank tests implemented using the survival (v3.6-4) R packages.

### Statistical analysis

Unpaired Student’s *t*-tests and Wilcoxon rank sum tests were used for comparing values between two groups. Fisher’s exact test was used for enrichment analyses. Survival analyses were conducted using Kaplan-Meier method with log-rank tests implemented in the survival packages in R. Benjamini-Hochberg was used to correct for multiple hypothesis testing and calculate FDR.

### Drug sensitivity data analysis

An unpaired Student’s *t*-test was used to calculate the statistical significance of area under the curve (AUC) for drug sensitivity data from Beat AML(25, 28). Benjamini-Hochberg approach was used to correct for multiple hypothesis testing and calculate FDR. Z-score normalized AUC values were calculated and multiplied by -1, to represent high values as sensitive to drug. AUC heatmap was created using the ComplexHeatmap package.

## Data availability

Published scRNA-seq datasets are available under the following GEO accessions: Murine HSC (GSE134242), normal human HSPC datasets (GSE75478(22) and GSE116256(10)), human AML datasets (GSE116256(10)), and murine LSC datasets (GSE140896(12)). The Zhang et al(24) datasets were provided by the authors. Newly preprocessed murine LSC raw count data are available on GitHub at https://github.com/yklee020/SCA. Beat AML RNAseq raw read count data and updated clinical data were downloaded from https://biodev.github.io/BeatAML2/. The TCGA LAML clinical data were obtained from the Genomic Data Commons (https://gdc.cancer.gov/about-data/publications/laml_2012), and the preprocessed RNA-seq raw read count data were acquired from(27). Sorted AML fraction RNAseq raw read count data and LSC activity annotations were obtained from GSE199452. Previously published microarray gene expression data were obtained from GSE6891 and GSE37642. Clinical data was obtained from Valk et al.(35) and from uploaded meta data in GEO. The results shown here are in whole or part based upon data generated by the TCGA Research Network: https://www.cancer.gov/tcga.

## Code availability

All R codes for this project and SCA package are available at https://github.com/yklee020/SCA.

## Results

### SCA identifies human HSPCs expressing the self-renewal gene expression program

SCA measures the similarity between a reference gene expression profile (the profile of interest) and the transcriptomic profile of individual cells within a query scRNA-seq dataset. This is achieved by computing the correlation coefficients of each cell to the reference profile, ranking the correlation values, and assessing the statistical significance of the assigned correlations. The reference profile is constructed using a marker gene-guided approach, such as differentially expressed genes. The reference profile can be derived from either scRNA-seq or bulk RNAseq datasets (**Figure 1**).

**Figure 1.**
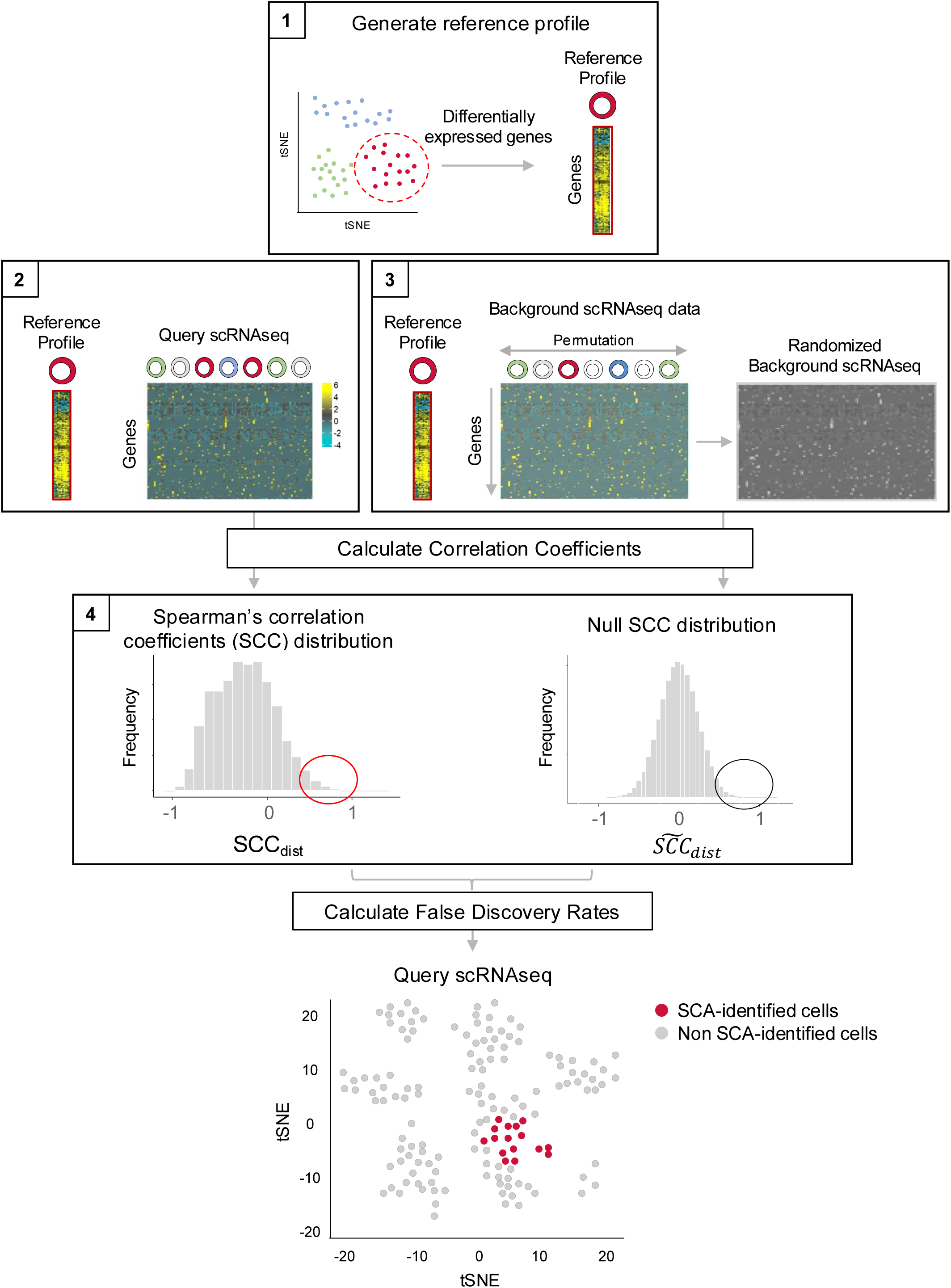
Schematic of Single cell Correlation Analysis (SCA). 1. First, a gene expression profile of interest, the reference profile, is generated through differential gene expression analysis of a reference dataset. **2.** Query dataset analysis: The similarity of the scGEP of each query cell to the reference profile, represented as a single vector of log-fold change values for genes, is calculated using a Spearman’s correlation coefficient (SCC). This generates observed SCC scores, SCC_dist_. **3.** Background data is randomized and a null distribution of SCC scores is generated between each pseudo cell in the randomized background dataset and the reference profile. **4.** Using the observed SCCs in the query dataset and the null distribution of the SCCs, 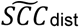. SCA estimates false discovery rate (FDR) for each query cell’s similarity score (each cell’s SCC). Cells having an SCC score that exceeds the SCC scores found in the null SCC distribution at a given FDR are defined as cells of interest that express the reference GEP.

To test SCA, we first used an *in vivo* validated, scRNA-seq signature of normal murine HSC self-renewal(*23*) as a reference profile and scRNA-seq datasets of normal human bone marrow precursors van Galen, 2019 #75;Velten, 2017 #231} for the query datasets (**Figure 2A**). In one of the query datasets(22), the self-renewal capacity of cells was experimentally validated using matched immunophenotypic marker expression and transcriptional data, along with functional self-renewal assays, providing matched functional and transcriptomic data. In the other query dataset, cell identities were defined transcriptionally through clustering and marker genes expression. We used these single cell datasets of HSC self-renewal to directly measure the accuracy, specificity, and precision of SCA in identifying self-renewing stem cells within the query dataset.

**Figure 2.**
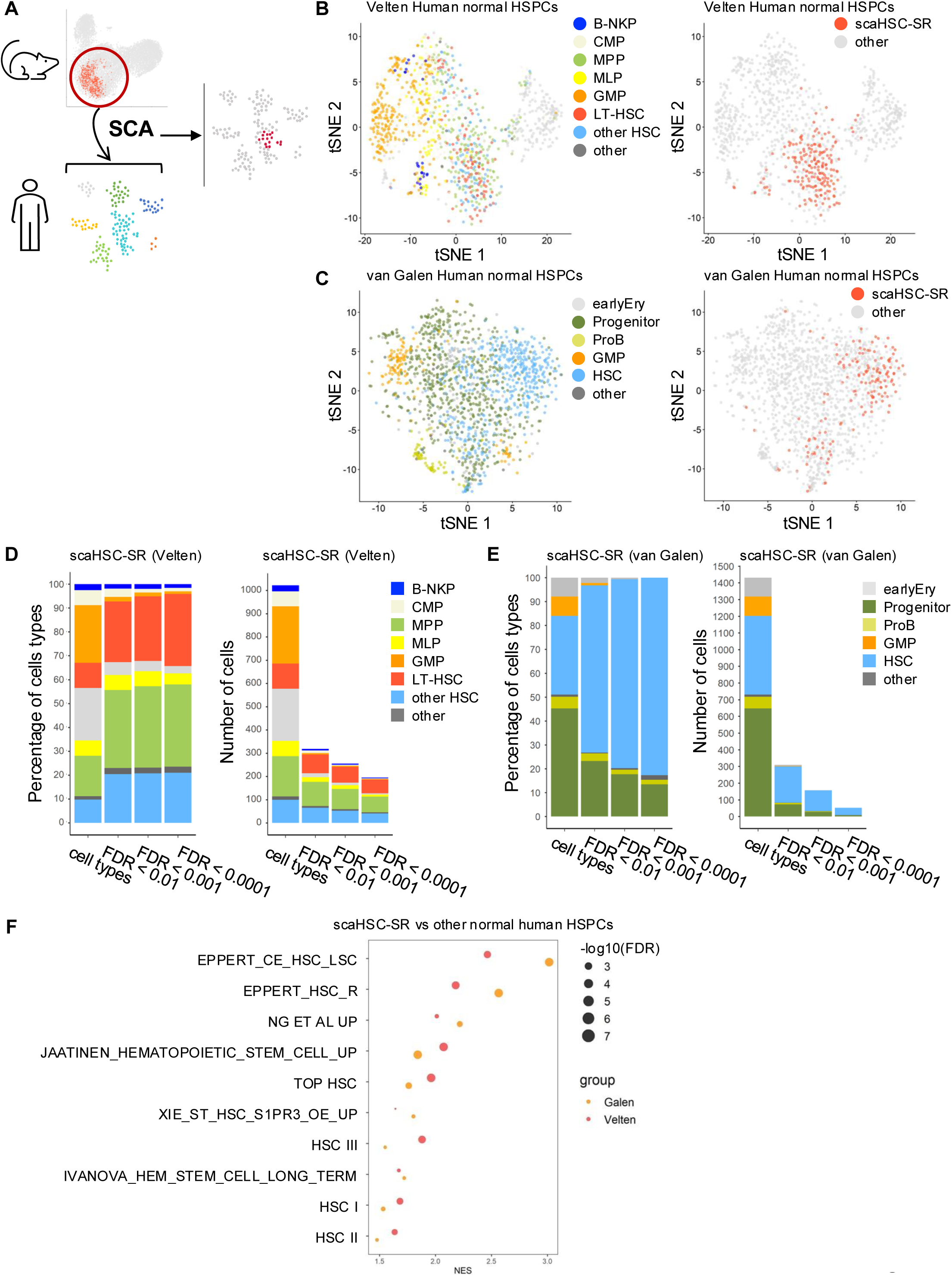
SCA identifies normal human hematopoietic stem cells that express the mouse hematopoietic stem cell self-renewal single cell gene expression profile. **A** Using an experimentally-validated mouse normal HSC self-renewal single cell gene expression profile as the reference profile(23), we applied SCA to query datasets of scRNA-seq of human normal HSPCs(10, 22). **B** tSNE visualization of the experimentally-validated, normal human HSPC dataset, Velten et al. HSPCs, is shown with the original cell type annotation provided by the study (left) and scaHSC-SR cells (SCA-identified cells that express mouse HSC self-renewal reference gene expression profile with FDR<0.0001, right). **C** tSNE visualization of a second independent normal human HSPC dataset, van Galen HSPC, is shown with the original cell type annotations provided by the study (left) and scaHSC-SR with FDR<0.001 (right). **D** Cell type annotation of the scaHSC-SR cells identified at different FDR cutoffs. **E** cell types of the scaHSC-SR cells identified at different FDR cutoffs. **F** GSEA was performed to compare scaHSC-SR with other HSPC cells in the scRNA-seq datasets. Each dataset was analyzed independently.

SCA computes a Spearman’s correlation coefficient (SCC) between the reference profile and each cell in the query datasets. To determine the range of SCC values that could be generated by randomized query data, SCA generates a null distribution of SCC scores using randomized data. Randomization of the query dataset is performed by permuting each gene’s expression value across cells to generate pseudo cells. A null SCC distribution was then generated by calculating SCCs between the reference profile and the pseudo cells in the randomized background data. The randomization process was repeated 1,000 times and the resulting null SCC distributions were incorporated into a comprehensive null distribution of SCC values. Finally, SCA estimated false discovery rate (FDR) for each cell’s similarity score, SCC, based on the empirical p-value derived from the null SCC distribution.

In both query datasets of human HSPCs, SCA identified cells expressing the reference self-renewal profile (scaHSC-SR) across varying FDR thresholds (**Figure 2B-E**). Since self-renewal capacity is limited to stem cells, we evaluated the cell type composition of scaHSC-SR cells in the both datasets (**Figure 2B-E**). In the Velten data, scaHSC-SR cells were primarily composed of true HSCs and progenitor cells, as defined functionally in that study (**Figure 2D**). In our analysis, we considered long-term hematopoietic stem cells (LT-HSCs) to be the cells that correspond to the mouse reference self-renewing HSCs and considered progenitor cells to be false positives. SCA assigns FDR value to each cell and allowing users to evaluate results across different levels of statistical stringency. As more stringent FDR thresholds were applied, the total number of scaHSC-SR cells decreased, but the proportion of LT-HSC among scaHSC-SR cells increased. This result reflects that more stringent FDR values provide higher statistical confidence and a reduced false positives rate. In the van Galen dataset, LT-HSC were not distinguished so we set HSCs as the cells of interest. Similar to our findings in the Velten analysis, the proportion of HSCs among scaHSC-SR cells increased with more stringent FDR thresholds (**Figure 2E**). Together, these results demonstrate that SCA can accurately identify the cells of interest with high precision. Importantly, we find that the false positive rate is minimized as the FDR cutoff becomes more stringent. Next, we performed gene set enrichment analysis (GSEA) to define biological pathways of scaHSC-SR cells in both datasets (**Figure 2F**). Each dataset was analyzed independently by comparing the scaHSC-SR cells to non-scHSC-SR cells. Among the pathways that showed concordant enrichment (at an FDR<0.05) in both datasets, we observed gene sets associated with normal HSC self-renewal, quiescent HSCs, and LSC self-renewal. These findings confirm that SCA can identify cells expressing transcriptional profiles of self-renewal.

### SCA outperforms existing scRNA-seq methods by optimizing false positive rate without compromising sensitivity or precision

Next, we compared SCA to several existing scRNA-seq analysis methods, scmap-cluster(17), SingleR(16), ScType(36), clustifyr(37), and AUCell(38). These methods can utilize a single reference profile vector to identify a cell type of interest. For these analyses, we constructed a common reference profile and applied it uniformly across all methods. Each method was then evaluated on the same human query datasets described above using multiple quality metrics. The performance of each method was assessed using the precision, sensitivity, F1 score (a harmonic mean of sensitivity and precision), and false positive rate (FPR) metrics. Additionally, we assessed how each method performed when using reference profiles of varying sizes by adjusting the number of differentially expressed genes (DEG) in the reference. First, we assessed inter-dataset classification performance by deriving the DEGs of the HSC population from one dataset (the van Galen HSPCs) and used that as reference profile to classify the Velten HSPCs as the query dataset. We selected the Velten dataset as the query dataset because the cell type annotations in this dataset are based on the measured immunophenotype. Notably, SCA precision was superior or equivalent to the other methods at all FDR cutoffs and reference profile sizes (**Figure 3A**). However, the sensitivity of SCA was significantly reduced when we used more stringent FDR values and the reference profile included a limited number of genes (**Figure 3B**). Therefore, SCA F1 scores dropped with more stringent FDR thresholds when the DEG list was small, but these SCA F1 scores remained high when the reference profile consisted of more DEGs (**Figure 3C**), indicating that a threshold of DEGs is important in the performance of SCA. Compared to other methods, SCA with permissive FDR thresholds (FDR<0.05 and FDR<0.01) achieved the highest F1 scores across most reference profiles sizes tested. When using larger reference profiles (1,000 and 2,000 DEGs), SCA exhibited comparable or superior F1 scores at all FDR threshold levels. Overall, SCA demonstrated higher precision with more stringent FDR cutoffs at the expense of sensitivity (**Figure 3A-B**).

**Figure 3.**
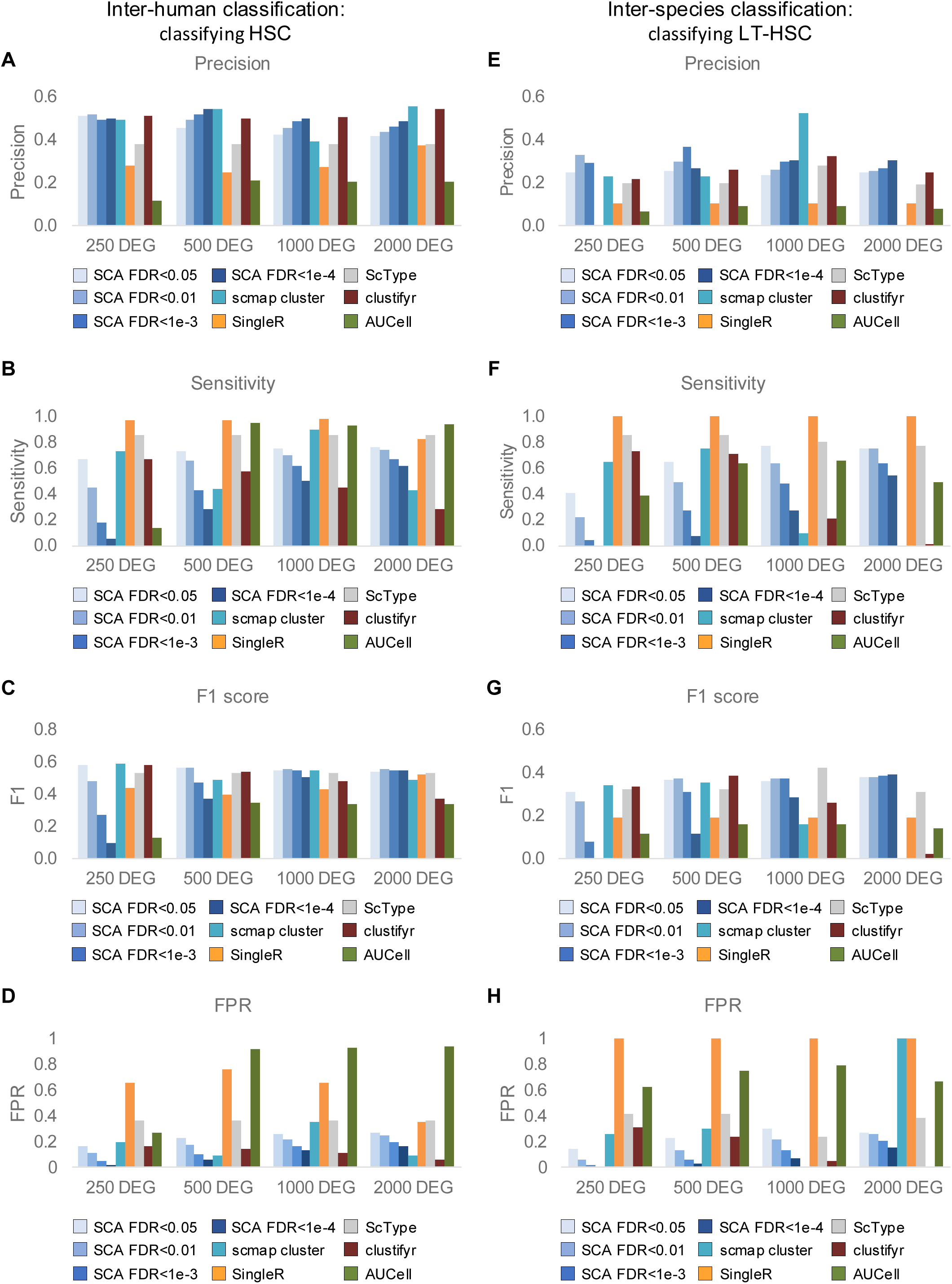
SCA outperforms existing scRNA-seq methods by optimizing false positive rates without compromising sensitivity and precision. Precision (**A, E**), sensitivity (**B, F**), F1 score (**C, G**), and FPR (**D, H**) were measured for both inter-human HSCs and inter-species classifications of LT-HSCs for SCA and other tools. The Velten data cell-type annotation is considered the gold-standard because these data are derived from index-sorted (i.e. flow cytometry-validated) cells. The van Galen data cell type annotation is based on transcriptomics clustering and marker gene. There are subtle differences in the annotation assignments across these two datasets. **A-D** Human HSC classification in the Velten human HSPC scRNA-seq dataset using HSC single cell gene expression profile from the van Galen normal human HSPC dataset as the reference profile. **E-H** LT-HSC classification in Velten HSPC scRNA-seq dataset, using mouse HSC self-renewal scRNA-seq data to generate the reference profile. Different numbers of differentially expressed genes were tested as reference profiles and the same reference profile was used in each analysis.

Next, we evaluated inter-species classification by using the mouse HSC self-renewal reference profile to classify human LT-HSCs. Consistent with the trends observed in the human dataset comparisons, F1 scores remained comparable or higher when larger reference profiles (1,000 and 2,000 DEGs) were used, and SCA consistently demonstrated superior precision compared to other methods (**Figure 3E and 3F**). As FDR cutoffs became more stringent, SCA maintained high precision while sensitivity decreased (**Figure 3E-F**). However, when smaller reference profiles were used, SCA showed lower sensitivities relative to other methods tested. In contrast, several of the other methods exhibited a decline in performance with larger reference profile sizes. Specifically, when using 1,000 or 2,000 DEGs, these methods showed reduced F1 scores, precision, and sensitivity compared to results obtained with smaller reference profiles. These findings highlight SCA’s robustness in handling larger reference profiles and its ability to maintain high precision across inter-species classification tasks.

In summary, SCA consistently achieves high precision across datasets, reference profile sizes, and FDR thresholds, and often outperforms existing methods in F1 score, particularly under permissive FDR settings. Larger reference profiles (1,000 and 2,000 DEGs) further higher F1 scores while maintaining precision across all levels of statistical thresholds and in both inter-dataset and inter-species comparisons. Collectively, these results demonstrate that SCA is comparable to and often superior in the performance of current reference-based cell-type classification methods.

One of the most challenging tasks for scRNA-seq analysis algorithms is achieving high specificity, in other words, minimizing false positive rates (FPR). Most existing methods are designed to optimize sensitivity. In contrast, SCA was designed to also optimize specificity by incorporating FDR calculation on the SCC. As demonstrated in the classification of HSCs and LT-HSCs (**Figure 3**), SCA exhibited decreasing FPR as the FDR thresholds became more stringent (**Figure 3D & 3H**). Compared to other methods, SCA consistently maintained a relatively low FPR while effectively balancing precision and sensitivity. In the case of human HSC classification, SCA, scmap-cluster and clustifyr achieved the lowest FPRs when larger reference profiles were used (**Figure 3D**). However, the low FPRs observed with scmap-cluster and clustifyr came at the cost of substantial reduction in sensitivity (**Figure 3B**), whereas SCA preserved high F1 scores and comparable sensitivity while minimizing FPR (**Figure 3B-D**). In the inter-species evaluation, where human query datasets were analyzed using murine HSC self-renewal reference profile, similar trends were observed. SCA, along with scmap-cluster and clustifyr, exhibited lower FPR with larger reference profiles (1000 and 2000 DEGs) (**Figure 3H**). Importantly, SCA retained higher F1 scores and sensitivity (**Figure 3G and 3F**), while maintaining low FPR. In contrast, scmap-cluster and clustifyr showed poor F1 scores and very low sensitivities. Additionally, SCA with permissive FDR thresholds (FDR < 0.05 and 0.01) achieved comparable or higher F1 scores while maintaining lower FPR than other methods. These results demonstrate that SCA offers a clear advantage in achieving superior or comparable classification performance by minimizing FPR without sacrificing sensitivity or F1 scores. Therefore, SCA is a novel, scRNA-seq analysis method with superior specificity without compromising precision or sensitivity. We next performed an intra-dataset assessment using a 5-fold cross-validation approach to evaluate each method’s generalization capabilities (**Supplemental Figure S1**). Similar to previous analyses, SCA achieved the lowest FPR under permissive FDR thresholds (FDR < 0.05 and 0.01), while still maintaining comparable F1 scores (**Supplemental Figure S1**).

### SCA can incorporate a common background dataset to allow comparisons of data across different experiments and sequencing technologies

Next, we determined whether SCA would permit direct comparisons across different datasets by incorporating a common background dataset. In previous analyses, the query dataset itself was randomized to generate the background dataset from which we derived the null SCC distribution. We evaluated whether a null SCC distribution derived from a common dataset, derived independently of the query dataset, could be used to analyze multiple query datasets. This approach would allow SCA to analyze multiple query datasets obtained from different experiments and capture technologies. By comparing observed SCCs in each query dataset to a common null SCC distribution, derived from a common background, results from different experiments could be directly compared. FDR calculated for SCA scores identified in each query dataset would thus be interpretable on a common scale, facilitating cross-experiment comparisons. To test this, we evaluated the reliability of using background datasets derived from other samples or sequencing technologies.

We first evaluated how the cell type diversity of the background dataset affects SCA results. To do this, we constructed multiple background datasets with different levels of cell type diversity. Using the van Galen and Velten datasets as background dataset, we subsetted the data by selecting cell populations to construct highly homogeneous background datasets (consisting of a single cell type) to increasingly heterogeneous background datasets (consisting of multiple cell types, **Supplemental Figure S2**). We included background datasets from two different studies in order to test the impact of different experiments and capture methods. This allowed us to test the reliability and generalizability of SCA when using background data from different sources and experimental conditions.

We measured the variance of data in each of these constructed background datasets to estimate cell type variability independently of cell type annotations. We initially assessed background dataset variability using principal component analysis (PCA, **Supplemental Figure S2A-B**). However, the variance captured by the higher variance in principal component (PC) did not consistently mirror the known biological gradient of cell type diversity. To address this limitation, we used the ROGUE statistic(30), an entropy-based metric that quantifies cell population purity based on gene expression, to evaluate the heterogeneity of each background dataset. The ROGUE statistic provided a purity metric that aligned with the biologically expected levels of diversity (**Supplemental Figure S2B**) decreasing as cell type diversity increased. Based on these results, we adopted the ROGUE purity scores as the metric to describe background dataset diversity in the following analyses.

We then evaluated how diversity of cell types in the background dataset, reflecting overall dataset heterogeneity, affects SCA classification performance. Specifically, we tested SCA in scenarios where the background dataset is highly homogenous, where most or all query cells resemble the reference profile. For example, we evaluated when both the background dataset and reference datasets are composed entirely of HSCs.

We measured F1 score and FPR in identifying cell of interest in this scenario. SCA was applied to identify Velten-derived LT-HSCs (**Figure 4**) and HSCs (**Supplemental Figure S3**) as the reference profiles and using the Velten HSPC dataset as the query dataset. The analysis utilized various background datasets, each constructed with progressively higher frequencies of the more diverse HSPC populations to vary cell type diversity (**Supplemental Figure S2A**). The results demonstrated that SCA failed or performed poorly in identifying LT-HSC and HSC and when the background datasets had high cell population purity, as indicated by ROGUE score > 0.94 (**Figure 4 & Supplemental Figure S3**). Similar observations were made in the van Galen HSPC dataset, where different subsets of HSPC populations were used as background data to identify HSCs (**Supplemental Figure S4**). In the Velten dataset, decreasing background purity (i.e., lower ROGUE scores) generally improved the identification of LT-HSCs. However, the dataset with the lowest purity (ROGUE score of 0.807) did not yield the best SCA performance (**Figure 4),** suggesting that overly heterogeneous backgrounds may also be suboptimal in certain cases. In contrast, for identifying broader HSC populations, increased background heterogeneity consistently enhanced SCA performance in both the Velten and van Galen datasets (**Supplemental Figure S3 and S4**). These findings suggest that SCA performs optimally when the background dataset contains high cell type diversity, with ROGUE scores below 0.94, within the tested conditions. This highlights the importance of selecting appropriately heterogeneous background dataset to maximize SCA’s classification accuracy.

**Figure 4.**
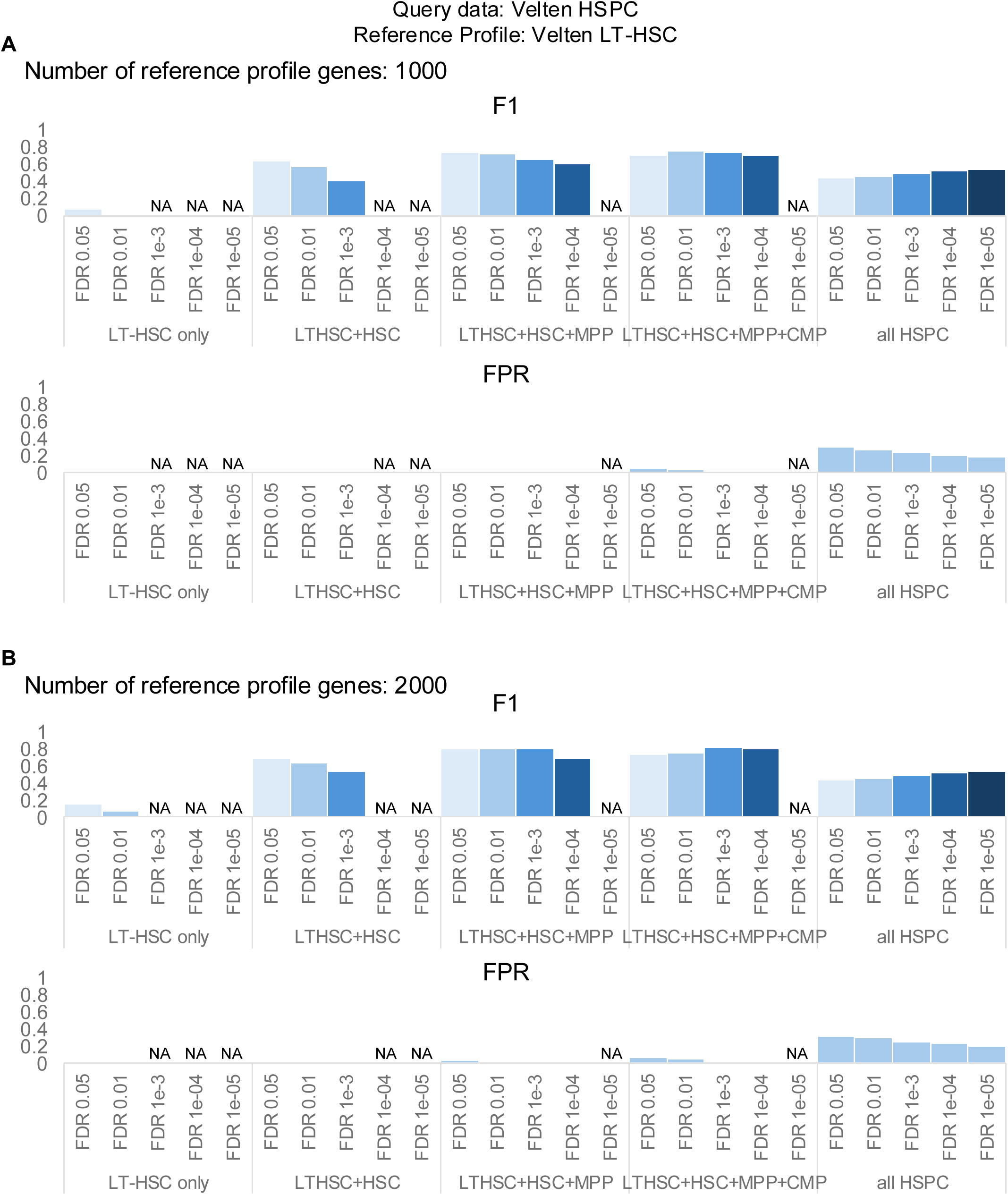
Incorporating background data with higher cell type diversity improves the SCA performance in identifying LT-HSCs in the Velten HSPC dataset. The impact of using different background datasets with varying levels of cell type diversity was evaluated. SCA was performed on Velten HSPCs as query data to identify LT-HSCs using the query data’s LT-HSCs as the reference profile with **A** 1000 genes and **B** 2000 genes. F1 scores and FPR were calculated for LT-HSC classification using different background datasets. Background datasets were created by sequentially adding cell populations from the query Velten data: LT-HSCs alone; LT-HSCs + other HSCs; LT-HSCs + other HSCs + MPPs; LT-HSCs + other HSCs + MPPs + CMPs; and all HSPCs. F1 scores and FPRs were measured at various FDR thresholds for each background dataset.

Next, we evaluated SCA performance using external background datasets derived from tissues with differing sample origins, single-cell capture technologies, and gene expression quantification methods. The Velten HSPC data was generated using Smart-seq2 protocol and read counts were measured for gene expression quantification(22), whereas the van Galen datasets were captured using Seq-well protocol with UMI counts (10). SCA was applied to identify LT-HSCs (**Supplemental Figure S5**) and HSCs (**Supplemental Figure S6**) in the Velten HSPC dataset, using the query data’s cell type of interest as the reference profiles. Analyses were performed with various background datasets, including the Velten HSPCs, van Galen unsorted BM, and van Galen HSPC (**Supplemental Figure S2A**). Despite differences in donor samples, capture technologies, and quantification methods, SCA consistently identified LT-HSCs and HSCs in the Velten dataset with high F1 scores and low FPR using the van Galen datasets as external background datasets (**Supplemental Figure S5 and S6**). Similarly, we also tested SCA on the van Galen normal HSPCs dataset as the query to identify HSCs, using its own HSC population as the reference profile. In this analysis, external background datasets were derived from different donors, capture technologies, and quantification methods (**Supplemental Figure S2A** and **Figure S7**). The van Galen unsorted BM data was derived from different donor sample from the query van Galen HSPC dataset(10) and the Velten data were generated using different donor sample, capture protocol and read count-based quantification. The results demonstrated that SCA performed better or comparable when using background datasets from different samples and technologies for identifying HSC populations (**Supplemental Figure S7**). Overall, these findings demonstrate that SCA can effectively utilize external background datasets from similar tissue origins, even when they differ in sample source, capture technology, or quantification methods. This finding demonstrates that SCA can be used to directly compare single cell transcriptional profiles across different experiments.

### Primary human AML scRNA-seq data harbors cells expressing experimentally validated self-renewal gene expression profile

We previously defined and experimentally validated single cell gene expression profiles of LSC self-renewal and proliferation in a murine model of AML(12). To determine whether human AML cells express these murine LSC self-renewal and proliferation gene expression profiles, we applied SCA to a publicly available scRNA-seq dataset of 16 adult human AML samples as the query dataset(van Galen(10), **Figure 5A**). The reference profiles consisted of murine LSC self-renewal and proliferation single cell gene expression profiles previously defined by our group(12). Using leukemia reconstitution and proliferation tracing studies and *in vivo* we previously demonstrated that self-renewing LSCs lack proliferation potential, while proliferating LSCs lack self-renewal capacity(12). Using SCA, we interrogated whether the van Galen dataset harbors cells that express these murine LSC self-renewal (scaLSC-SR) or LSC proliferation (scaLSC-Pro) profiles (**Figure 5A-B**). All 16 human samples were used as background data for SCA FDR estimation. Our findings revealed that all 16 adult AML samples harbor scaLSC-SR cells (FDR<1x10^-5^). In contrast, half of the samples harbor scaLSC-Pro cells (FDR<1x10^-5^).

**Figure 5.**
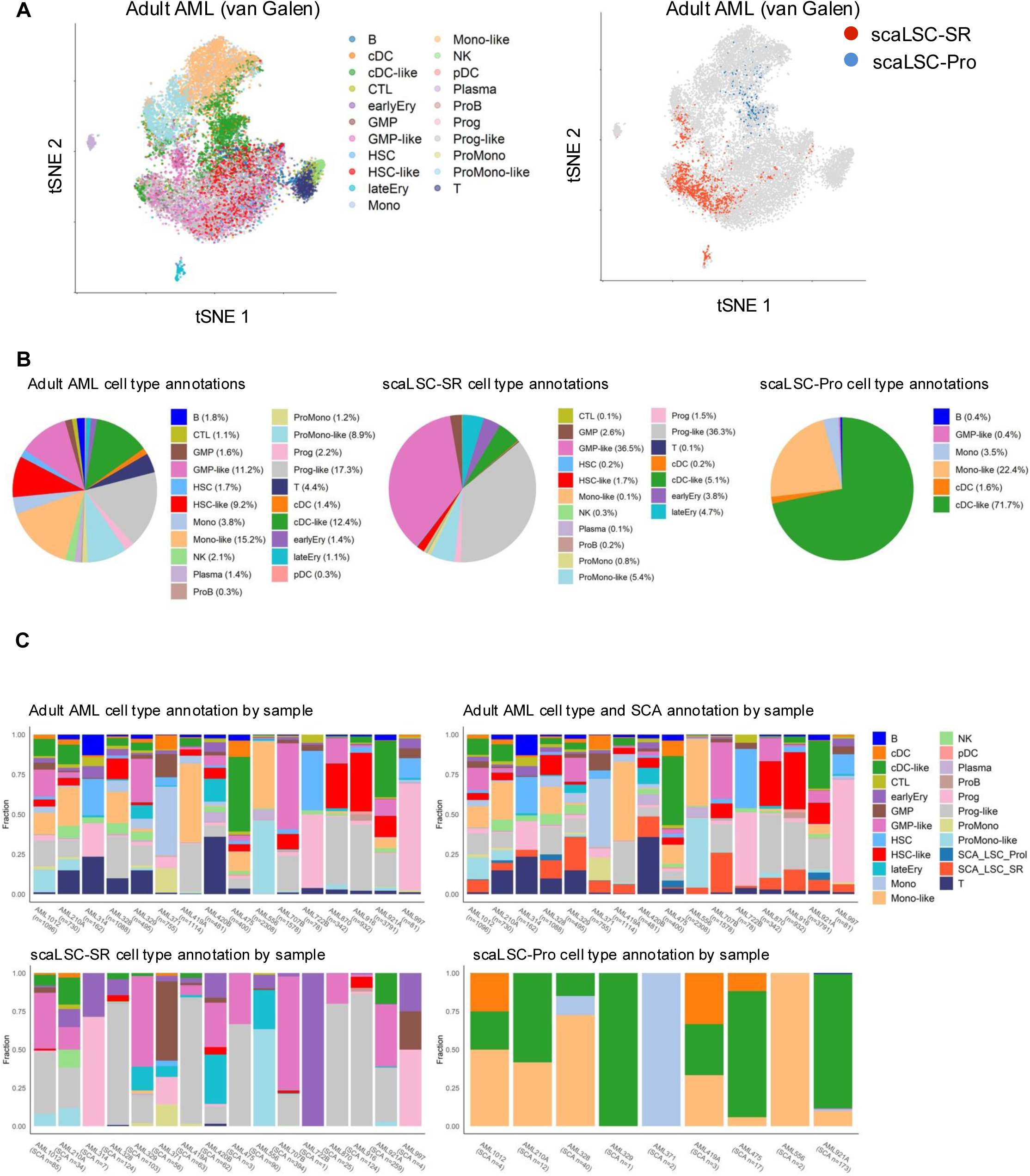
SCA identifies adult AML cells expressing mouse LSC self-renewal and proliferation single cell gene expression profiles. **A** tSNE visualization of 16 adult human AML samples’ scRNA-seq data is shown with the cell type annotations provided in the study[ref: van Galen] (left) and with SCA-identified AML cells expressing the reference mouse LSC self-renewal (scaLSC-SR, red) and LSC proliferation (scaLSC-Pro, blue) single cell gene expression profiles with FDR<1x10^-5^. **B** Pie charts display cell type compositions in all samples in the dataset (left), within scaLSC-SR cells (middle), and within scaLSC-Prol cells (right). The percentages of each cell type are included in the graph legends. **C** Bar plots showing original cell type compositions within each AML sample (top left). Fractions of cell types are displayed for each sample; along with fractions of scaLSC-SR and scaLSC-Pro cells in each sample (top right). Additional plots show cell type fractions within scaLSC-SR (bottom left) and within scaLSC-Pro (bottom right) across samples.

We then examined the cell type composition of scaLSC-SR and scaLSC-Pro cells based on the previously defined cell type annotation from the van Galen study(10). In that study, AML bone marrow cells were classified as normal or malignant with malignant cells labeled according to their closest normal counterpart (e.g., “-like” cells). Among the scaLSC-SR population from 16 adult samples, Progenitor-like (36.3%) and GMP-like (36.5%) cells represented the majority of the cell compositions (**Figure 5B**). This finding suggests that the scaLSC-SR closely resembles human malignant progenitor cells, which are thought to harbor functional LSCs. On the other hand, scaLSC-Pro cells were primarily composed of cDC-like (71.7%) and Mono-like (22.4%) cells, indicating scaLSC-Pro cells closely resemble more mature malignant populations lacking self-renewal capacity. Additionally, the composition of cell types within scaLSC-SR and scaLSC-Pro varied across individual samples, consistent with existing literature reports LSCs abundance variability between patients(6, 8)(**Figure 5C**). These findings support the utility of SCA in identifying transcriptionally distinct subpopulations associated with LSC self-renewal and proliferation across AML samples.

Using GSEA, we compared the biological pathways of scaLSC-SR cells to all other AML cells in the data. This analysis revealed that scaLSC-SR cells were significantly enriched in gene sets associated with normal hematopoietic stem and progenitor cells, LSC self-renewal, and oxidative phosphorylation (OXPHOS; FDR < 0.05, **Supplemental Figure S8A**). These pathways align with known molecular features of LSC self-renewal. Additionally, GSEA results indicated significant depletion of gene sets related to apoptosis, hematopoietic differentiation, KRAS signaling, allograft rejection, and inflammatory response. These findings are consistent with prior studies showing that LSCs are resistant to apoptosis and can evade immune responses, both critical features that contribute to their persistence and therapy resistance(3, 4). To further characterize the molecular features of self-renewing LSCs, we examined the expression of known LSC marker proteins, typically identified in CD34+CD38-populations, across different cell types. Using cell surface marker-encoding genes as surrogates, we performed differential gene expression analysis across 16 AML samples. This analysis identified cell surface marker-encoding genes significantly upregulated in scaLSC-SR cells (**Supplemental Figure S8B**). Among these, we identified CD34, a well-established LSC marker(3, 6, 8), and CD38, which is often expressed in a more differentiated progenitor compartment(3, 6, 8) were differentially upregulated in scaLSC-SR cells.

Furthermore, both CD96 and CD200, previously described as immunophenotypic markers of functional LSCs(39, 40), were also upregulated in scaLSC-SR cells. These findings suggest that scaLSC-SR cells express classical LSC markers but also exhibit transcriptional features consistent with functional LSCs. The additional genes and pathways identified in these cells provide further insights into the molecular underpinnings of LSC self-renewal and differentiation states.

Next, we applied SCA to a scRNA-seq dataset of pediatric AML (24) to determine whether the murine reference profile for LSC self-renewal and proliferation could be detected in this population. This dataset includes scRNA-seq data of 13 pediatric AML samples. All 13 pediatric AML samples were pooled to generate the background data for SCA. SCA revealed that all 13 pediatric AML samples harbor both scaLSC-SR and scaLSC-Pro cells (with an FDR<1x10^-6^, **Figure 6A**). These findings indicate that, similar to adult AML, pediatric AML harbors cells expressing the murine LSC self-renewal and proliferation gene expression profiles (scaLSC-SR and scaLSC-Pro cells).

**Figure 6.**
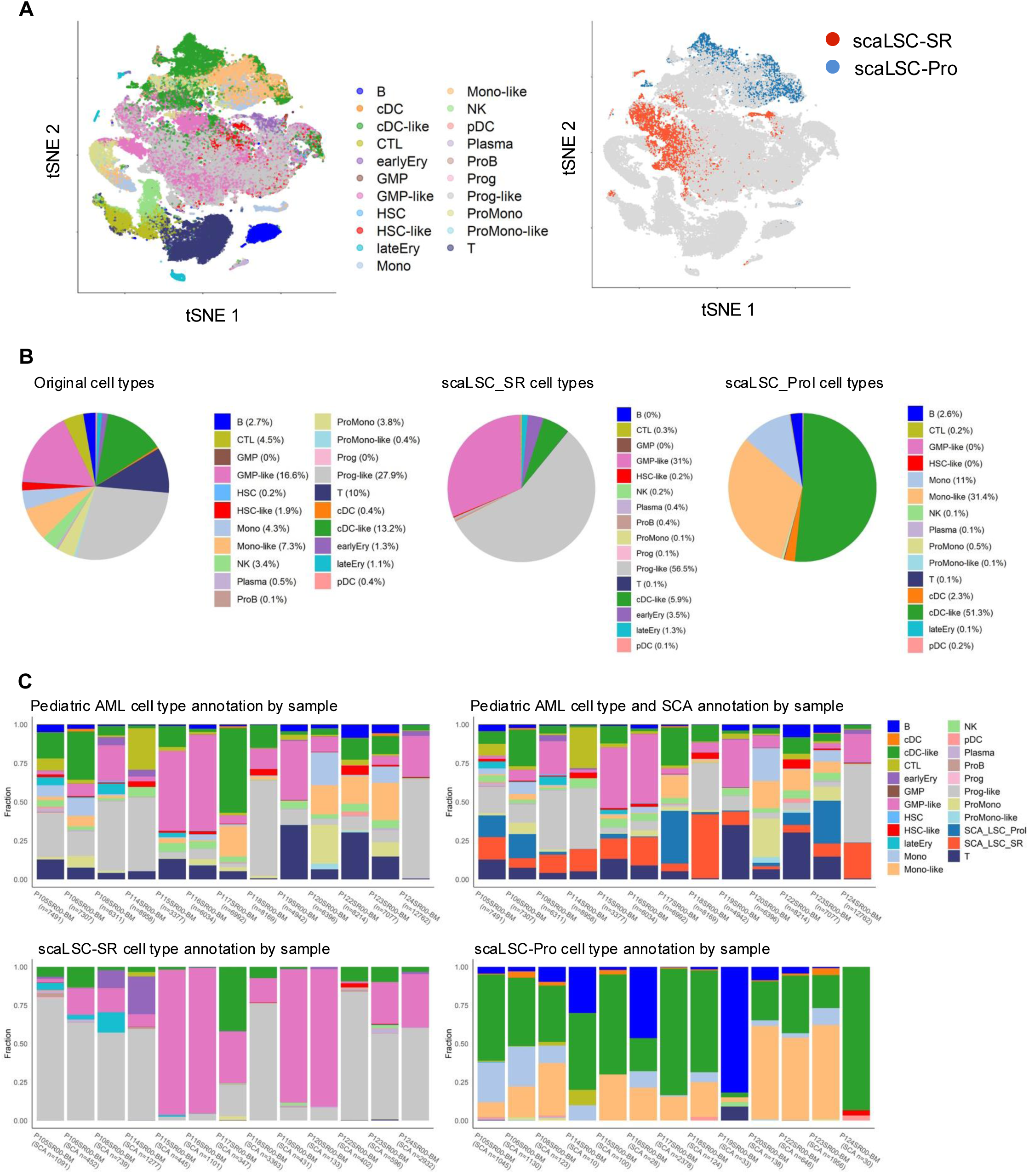
SCA identifies pediatric AML cells expressing the mouse LSC self- renewal and proliferation single cell gene expression profiles. **A** tSNE visualization of 13 pediatric human AML samples’ scRNA-seq data (Zhang et al. Genome Biology 2022) is shown. Cell type annotation was transferred to the pediatric dataset using the AML scRNA-seq data from van Galen et al 2019 using SingleR (left). SCA-identified AML cells expressing the mouse LSC self-renewal (scaLSC-SR, red) and LSC proliferation (scaLSC-Pro, blue) single cell gene expression profiles with FDR< 1x10^-6^ (right). **B** Pie charts display cell type compositions in all samples in the dataset (left), within scaLSC-SR cells (middle), and within scaLSC-Prol cells (right). The percentages of each cell type are included in the graph legends. **C** Bar plots showing cell type compositions within each AML sample. Fractions of cell types are displayed for each sample (top left); along with fractions of scaLSC-SR and scaLSC-Pro cells in each sample (top right). The lower plots show cell type fractions within scaLSC-SR (bottom left) and within scaLSC-Pro (bottom right) across samples.

To compare the cell type compositions of the scaLSC-SR and scaLSC-Pro cells in the pediatric and adult AML samples, we first standardized cell type annotations across both datasets. We used the cell type annotations defined in the van Galen, which distinguishes malignant AML cells from normal bone marrow cells using single cell genotyping and gene expression profiling. This annotation was applied to the pediatric AML dataset using SingleR(16). In pediatric samples, scaLSC-SR cells were predominantly Progenitor-like (56.5%) and GMP-like (31.0%), mirroring the distribution observed in adult AML (**Figure 5B** & **Figure 6B**). Similarly, cDC-like (51.3%) and Mono-like (31.4%) cells were the major constituents of scaLSC-Pro in pediatric samples (**Figure 6B**), consistent with adult AML (**Figure 5B**). The frequencies of scaLSC-SR and scaLSC-Pro cells varied across individual pediatric patients (**Figure 6C**), as observed in adult patients (**Figure 5C**).

To further investigate the biological features of scaLSC-SR cells, we performed GSEA comparing scaLSC-SR cells to all other AML cells in both adult and pediatric datasets. This analysis showed concordant enrichment of gene sets associated with normal hematopoietic stem and progenitor cells, LSCs self-renewal, and OXPHOS in both adult and pediatric scaLSC-SR population (**Supplemental Figure S9A**). These pathways are characteristic features of LSC self-renewal and suggest shared molecular pathways between adult and pediatric scaLSC-SR cells. GSEA also showed significant depletion of gene sets related to apoptosis, hematopoietic differentiation, KRAS signaling, allograft rejection and inflammatory response in both adult and pediatric datasets. These findings are consistent with prior studies showing that LSCs are resistant to apoptosis and can evade immune responses(3, 4). The downregulation of differentiation pathways also described the stem-like features of scaLSC-SR cells. We also identified cell surface marker-encoding genes that were significantly upregulated in pediatric scaLSC-SR (**Supplemental Figure S9B**). Many of these markers overlapped with those found in adult scaLSC-SR, while some were unique to the pediatric scaLSC-SR cells. The shared upregulation markers included CD34, CD38, CD200, CD320, and CD96 between adult and pediatric scaLSC-SR. These markers are associated with stemness and have been implicated in LSC biology(3, 6, 8, 39, 40). We also identified the cell surface marker-encoding genes uniquely upregulated in pediatric scaLSC-SR cells, such as CD69. Previous study found that CD69 is highly expressed in a chemoresistant HSC-like population in pediatric AML. These differences may reflect potential differences in the molecular characteristics of pediatric and adult LSCs.

### LSC Self-renewing cell composition is associated with mutational status in AML

Previous studies have demonstrated that AML cell type compositions vary across genetic subgroups of AML and are associated with poor outcomes(9, 25). To investigate whether the abundance of scaLSC-SR cells is associated with specific mutational patterns in AML, we performed gene expression deconvolution analysis to estimate cell type compositions within bulk AML gene expression profiles(25–27). For this analysis, we used cell type annotations established by van Galen et al, which distinguish normal bone marrow from malignant AML cells, and incorporated our reference scaLSC-SR and scaLSC-Pro cell types. Using CIBERSORTx(33), we deconvoluted the Beat AML and TCGA LAML datasets (**Supplemental Figure S10A**), quantifying the relative frequencies of scaLSC-SR compositions across samples.

We next examined the association between scaLSC-SR composition and mutational status in AML (**Supplemental Figure S10B**). For each of the most frequently observed mutations in AML, we compared scaLSC-SR compositions between mutant and wild-type samples (**Figure 7**). In both the Beat AML and TCGA LAML datasets, samples harboring *TP53Mut* and *NRASMut* exhibited significantly higher scaLSC-SR compositions compared to wild-type samples. These findings align with prior studies demonstrating that both mutations enhance self-renewal capacity in AML(12, 13, 41, 42). Conversely, samples with *NPM1Mut* and *FLT3-ITDMut* showed significantly lower scaLSC-SR frequencies in both datasets compared to wild-type samples. This observation is consistent with *NPM1* mutation in AML is classified as a favorable risk in the absence of additional high-risk mutations(43). In the Beat AML dataset, additional *NF1Mut* and *KRASMut* mutations were associated with higher scaLSC-SR compositions, while *TET2Mut* and *IDH2Mut* were associated with lower scaLSC-SR frequencies. Similarly, in the TCGA LAML dataset, *FLT3Mut* were also associated with reduced scaLSC-SR composition. These results suggest that the abundance of scaLSC-SR populations varies across genetic subtypes in AML, implicating the influence of specific mutations on self-renewal capacity and cellular composition.

**Figure 7.**
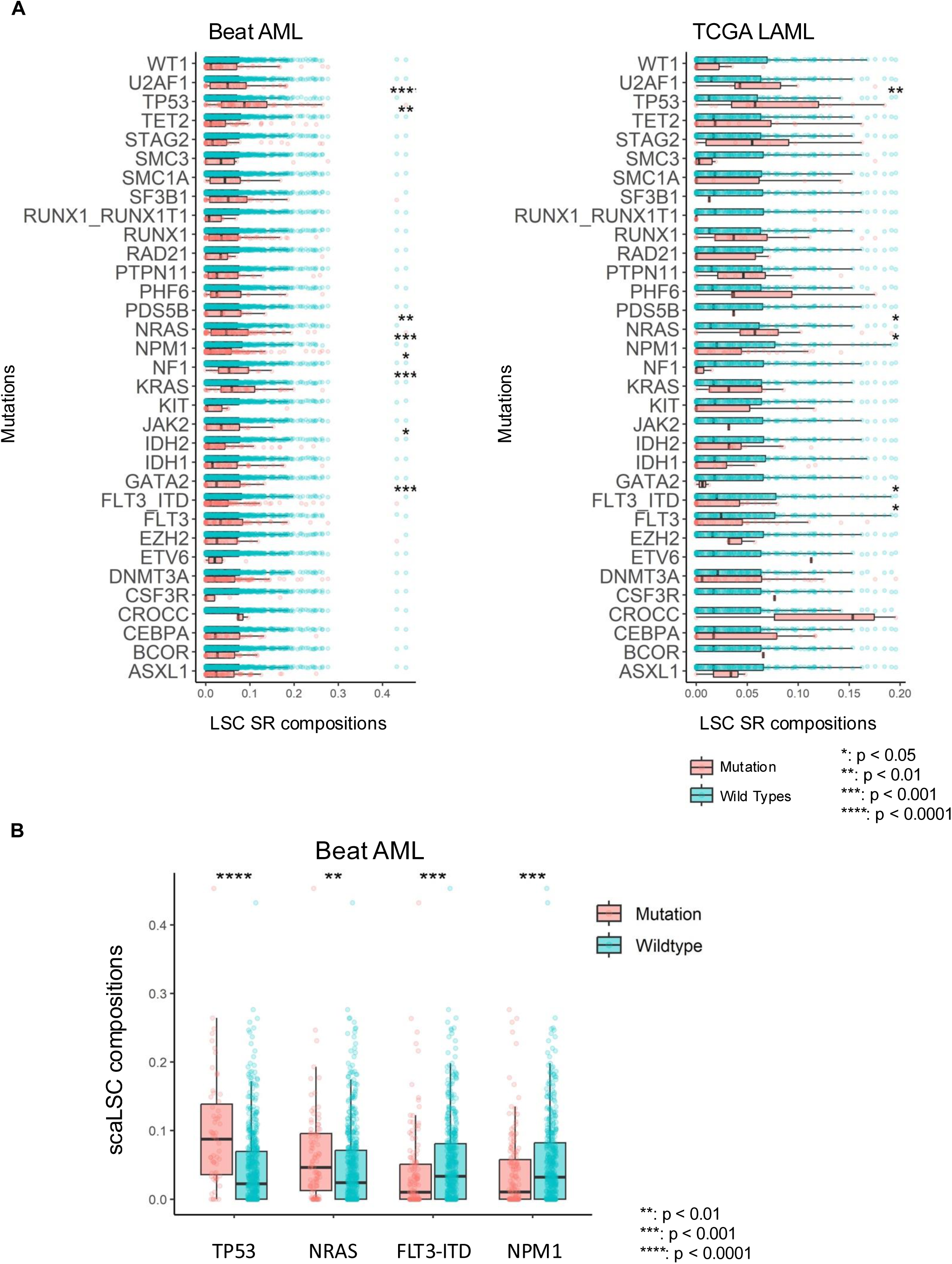
Different AML mutational subtypes show distinct scaLSC-SR compositions. **A** Enrichment of scaLSC-SR cell type was compared between the mutation and wild type groups for each of the most frequently mutated genes in Beat AML and TCGA LAML datasets. **B** Mutational subgroups that were concordantly enriched (*TP53Mut* and *NRASMut*) or depleted (*FLT3-ITDMut* and *NPM1Mut*) for LSC-SR compositions with statistically significant associattions in both Beat AML and TCGA LAML are illustrated. Relative abundances of scaLSC-SR cell type were compared using a two-sided Wilcoxon rank-sum test (p-values are presented for each comparison).

### scaLSC-SR gene signature scores are associated with AML outcomes

We further tested whether the scaLSC-SR gene signature is associated with AML patient outcomes using bulk gene expression profiles from four independent AML cohorts (**Figure 8A**). First, we defined a human-specific scaLSC-SR gene list by selecting upregulated DEG (with fold change > 2 and FDR < 0.05) identified in the van Galen AML scRNA-seq dataset(10). We applied a Cox proportional hazards model with LASSO regularization (LASSO Cox regression) and trained on the GSE6891 dataset, which consists of 503 newly diagnosed AML patient samples. This process identified 28 genes signature, we termed LSC-SR28, whose weighted expression sum was used to compute an LSC-SR28 score for each patient.

**Figure 8.**
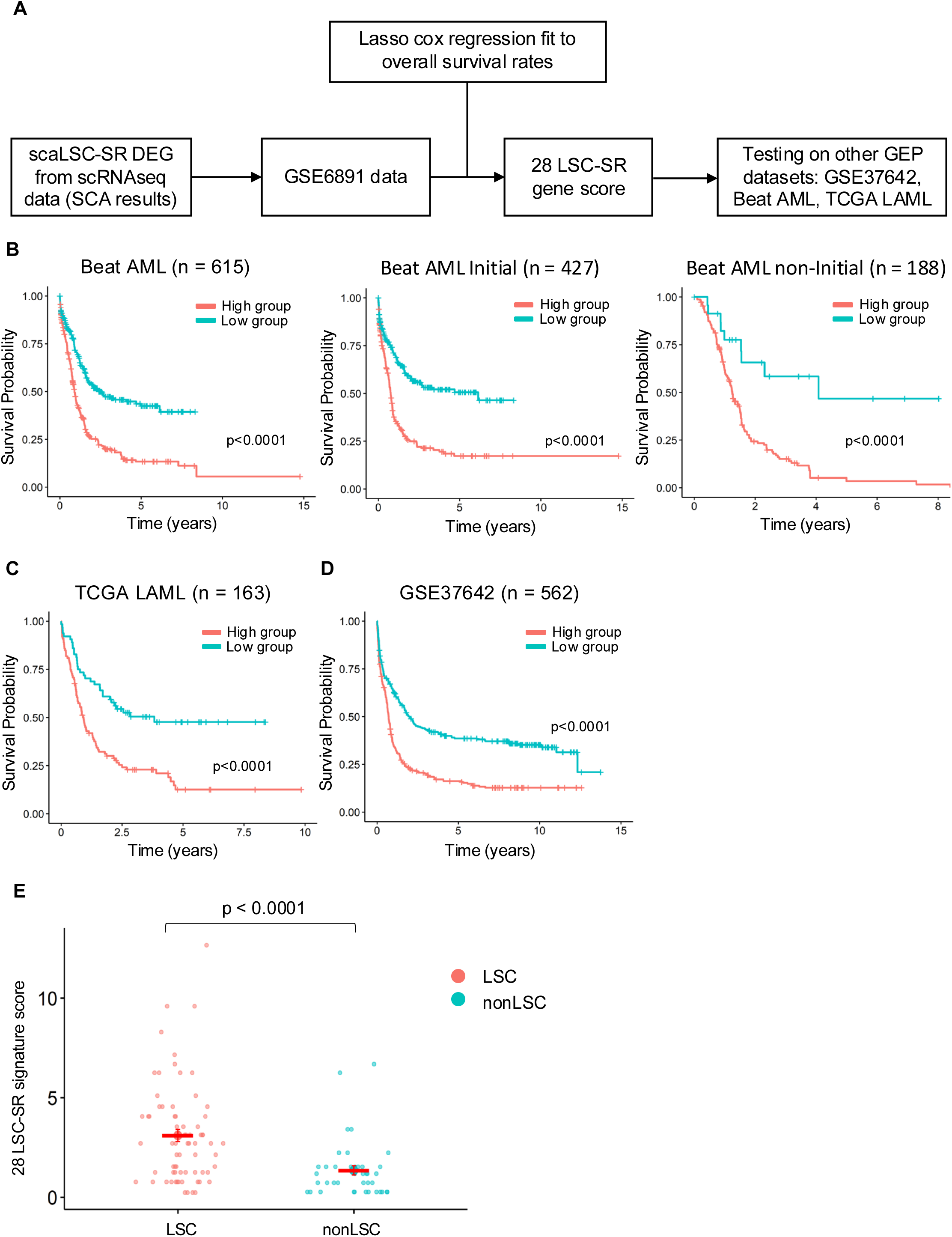
High scaLSC-SR gene signature scores are associated with overall survival in intendent AML cohorts. **A** Schematic illustrating the generation of the scaLSC-SR gene signature and its testing using gene expression profiles and overall survival rates in three human AML datasets. **B** Kaplan-Meier curves comparing estimates of overall survival (OS) between high and low scaLSC-SR signature score groups in the Beat AML (all samples, n = 615), Beat AML initial diagnosis samples (n = 427), Beat AML non-initial diagnosis samples (n = 188). **C** Kaplan-Meier curve comparing OS in TCGA LAML (all initial diagnosis samples, n = 163) and **D** a German dataset from the AMLCG 1999 trial, GSE37642 (all diagnosis samples, n = 562). P-values were calculated using the log-rank test. **E** scaLSC-SR gene signature scores were calculated in 111 bulk RNAseq samples from sorted AML fractions previously tested for LSC self-renewal activity in Ng 2016. scaLSC-SR gene signature scores between LSC (engrafted) and nonLSC (non-engrafted) groups were compared using a two-sided Wilcoxon rank-sum test.

We evaluated the prognostic relevance of the LSC-SR28 gene signature across three independent AML cohorts (GSE37641, Beat AML and TCGA LAML). Patients were stratified into high- and low-risk groups based on an LSC-SR28 score cutoff that maximized log-rank test values for overall survival (OS) differences. In all datasets, including Beat AML (n = 615 AML samples from diagnosis and later stage in the disease course), diagnostic Beat AML cases (n = 427), TCGA LAML (diagnostic cases, n = 163), and GSE37642 (German AMLCG 1999 trial, n = 562 diagnostic cases), patients with high LSC-SR28 scores consistently had significantly poorer OS in each dataset (**Figure 8B-D**). To further validate the association between LSC-SR28 scores with leukemic self-renewal, we calculated the scores in a bulk RNAseq dataset of sorted AML fractions with experimentally validated LSC self-renewal function(8, 9). LSC-SR28 scores were significantly higher in LSC-enriched fractions compared to non-LSC fractions (p < 0.0001), demonstrating that the LSC-SR28 signature reflects self-renewal activity in human AML (**Figure 8E**). Overall, we found that high LSC-SR28 scores were strongly associated with both poor OS and experimentally validated functional LSC self-renewal in AML. This highlights the potential utility of LSC-SR28 signature for risk stratification and therapeutic targeting in AML.

We further investigated the mutational landscape of patients with high LSC-SR28 scores in the Beat AML and TCGA LAML datasets (**Supplemental Figure S11A-C**). In both datasets, high scoring patients exhibited significant enrichment of *TP53Mut* and depletion of *RUNX1-RUNX1T1* fusions (**Supplemental Figure S11D**). In the Beat AML dataset, high LSC-SR28 scores were additionally associated with *DNMT3AMut*, *FLT3-ITDMut*, *JAK2Mut, RUNX1Mut, SF3B1Mut, TP53Mut*, as well as depletion of biallelic *CEBPAMut*. While the TCGA LAML dataset displayed similar trends, several mutation associations did not reach statistical significance, likely due to the smaller sample size. Overall, we find that high LSC-SR28 score patients showed an enrichment of genetic features that define poor risk AML and a depletion of favorable risk features. This further supports the clinical relevance of the LSC-SR28 signature in stratifying AML patients.

### High LSC-SR28 score AMLs show unique drug resistance profile

We tested whether AML samples with high LSC-SR28 scores exhibit distinct drug resistance profiles, we analyzed *ex vivo* drug response data from the Beat AML dataset, which includes response to 122 small molecule inhibitors. The high LSC-SR28 score group samples showed a significantly different drug sensitivity pattern compared to the low-score group (**Figure 9A**). Further analysis revealed that high LSC-SR28 score samples were sensitive to several small molecule inhibitors, including Venetoclax, a BCL-2 inhibitor that is part of the current standard of care for AML therapy (**Figure 9B-C**). This finding aligns with previous reports indicating that LSCs are particularly sensitive to Venetoclax treatment(5), highlighting that the patients with high LSC-SR28 scores may benefit from targeted therapies that aims for LSC vulnerabilities.

**Figure 9.**
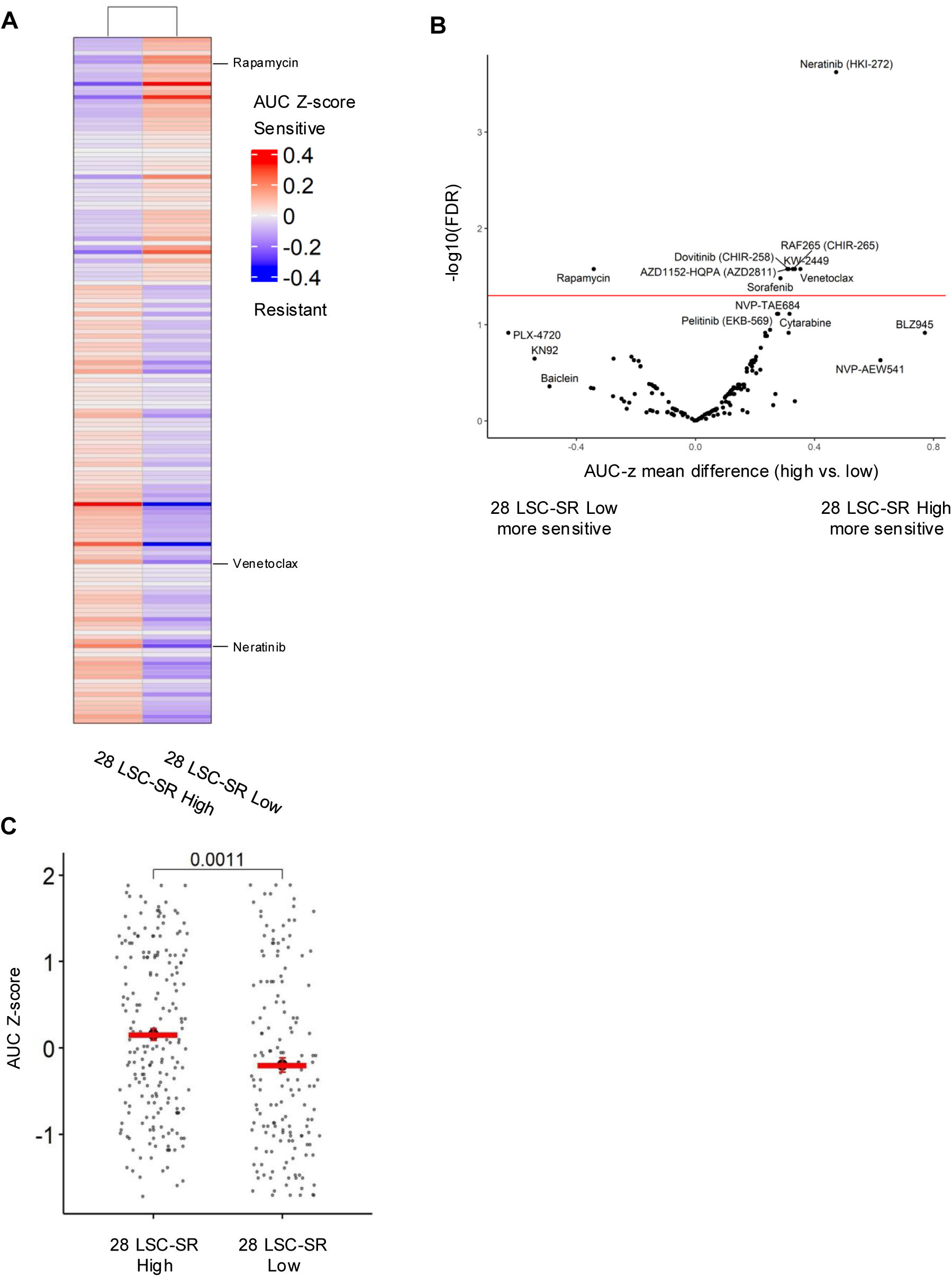
AML samples with high scaLSC-SR scores exhibit unique drug-resistant patterns and sensitivity to Venetoclax. *Ex vivo* drug sensitivity data generated from 122 small molecule inhibitors in the Beat AML dataset. Area under the curve (AUC) values were Z-score transformed and multiplied by -1 to generate AUC Z-scores. High Z-score indicates drug sensitivity. **A** Heatmap of the entire dataset. **B** Unpaired Student *t*-test was used to compare the average differences in AUC drug responses between scaLSC-SR score high versus scaLSC-SR signature low samples. Multiple hypothesis testing was corrected using the Benjamini-Hochberg method to calculate FDR. Red line indicates cutoff of FDR=0.05. **C** Venetoclax sensitivity in scaLSC-SR signature high and low samples. An unpaired Student *t*-test was used to compare the average AUC differences between scaLSC-SR signature high to low samples.

## Discussion

SCA provides a robust and statistically principled framework for identifying rare and biologically meaningful cell populations across heterogeneous single-cell datasets. By utilizing permutation-based FDR estimation, SCA confidently determines whether a query scRNA-seq dataset contains cells expressing a gene expression program of interest. The estimated FDR provides a standardized thresholds that effectively balances low false positives rates and high sensitivity in detecting rare subsets in complex tissues. In our case, we used SCA to demonstrate that human AML samples harbor cells that express a functionally validated gene expression profile of self-renewal. Compared with existing reference-based annotation tools, SCA achieves superior or comparable accuracy and sensitivity while maintaining low false positive rates across diverse datasets in identifying rare cell types, including identification of LT-HSCs and HSCs in normal HSPCs. In addition, SCA is designed to offer significant flexibility, allowing the use of sorted bulk RNA-seq data or any ranked gene list as a reference profile to query scRNA-seq datasets. Importantly, SCA provides statistical assessments of its results and can be used to directly compare independent datasets Together, these features make SCA a unique tool for rigorous annotation and analysis of single-cell transcriptomic data.

One novel benefit of SCA is its ability to directly compare cells across independent datasets. SCA utilizes a rank correlation coefficient and external background datasets to estimate common FDR thresholds, providing statistically consistent criteria for evaluating SCA results across samples. In general, many approaches compare cell type annotation across different datasets through data integration(44–46) which align datasets into the shared embedding space but these approaches rely on arbitrary or relative thresholds when assigning cell-type annotations. Similarly, supervised cell type annotation algorithms (16, 17, 36–38) compute similarity, signature, or probabilistic scores for each cell and then set relative cutoffs based on score distributions specific to individual dataset. Methods like AUCell also determine whether each cell upregulates gene-set level GEP using intra-sample thresholds. Other methods assign cell types using the highest similarity scored cell type(16, 17, 36–38) or voting among highest scoring cells (17, 20, 37). However, their thresholding strategies lack a statistical framework, such that the thresholds they assign are arbitrary. In contrast, SCA overcomes the use of arbitrary or relative classifications by introducing a permutation-based FDR framework that provides standardized, and statistically-defined thresholds.

In our work, we quantitatively evaluated how different levels of cell type diversity in reference scRNA-seq data impacts SCA performance. We utilized previously developed cell diversity statistics, the ROGUE statistic(30), to measure cell type diversity and show that higher heterogeneity within the reference dataset enhances SCA’s sensitivity and robustness in detecting HSC programs in normal human bone marrow scRNA-seq data.

Most computational tools that were designed to transfer cell annotations utilizing large-scale well-annotated reference dataset emphasize the choice of reference data is critical for the cell type annotation performance. However, the minimal or optimal level of cell type diversity required for accurate classification is not commonly reported or systematically evaluated. Most studies have focused on the effects of unmatched cell types between reference and query datasets or on the impact of cell-number imbalance among cell types within the reference. For example, PopV (47) reported that annotation accuracy declines when the reference lacks specific cell types present in the query.

While deCS(48) reported similar results; this study improved performance by combining cell type marker genes from multiple and different tissue type references. Likewise, other studies(49, 50) noted that unequal cell numbers in each cell type in reference datasets impact classification results. Similar to our study, Ma et al(51) demonstrated that combining multiple reference samples from different conditions improved supervised cell type annotation performance, independent of the increase in total cell numbers within the reference, suggesting that the improvement was primarily driven by increased biological variability. However, these studies did not quantitatively assess how variation in reference cell-type composition contributes to these improvements. Here, we demonstrate that cell type diversity in reference data is essential for SCA’s performance and quantify the level of diversity for optimal performance.

Adult and pediatric AML differ in their mutational landscapes, chromosomal abnormalities, and other molecular features (43, 52). Yet, the functional and transcriptional features of human LSCs have not been extensively compared between adult and pediatric disease. Our analysis revealed that both adult and pediatric AML harbor malignant cells that express a conserved LSC self-renewal program. Previous studies investigated the mechanism of LSC self-renewal using GEPs of functionally validated LSC-enriched compartments in adult AML(6, 8). The LSC gene signature, developed from these adult bulk GEP datasets, has demonstrated strong prognostic value in both adult and pediatric AML(8). Subsequent studies have validated this LSC gene signature in independent pediatric cohorts, confirming that the stemness program associated with LSCs is both prognostically relevant and largely conserved across different age groups (53, 54). Despite this cross-cohort validation, a pediatric-specific functional LSC GEP has not yet been characterized. Using SCA, we identified scaLSC-SR cells across multiple adult and pediatric AML patient samples and demonstrated that the transcriptional program of LSC self-renewal is shared across different age groups.

These scaLSC-SR cells predominantly consisted of Prog-like and GMP-like AML cells in both age groups. Moreover, both adult and pediatric scaLSC-SR cells shared enrichment in pathways associated with LSC self-renewal, normal hematopoietic stem and progenitor cells, and oxidative phosphorylation, all of which are hallmarks of LSC biology(5, 55). These cells were also depleted in pathways linked to apoptosis, differentiation, and immune responses, consistent with the known ability of LSCs to resist cell death and evade immune surveillance, enabling their persistence and therapy resistance. Notably, scaLSC-SR cells in both adult and pediatric AML consistently upregulated key cell surface marker-encoding genes such as CD34, CD38, CD200, CD320, and CD96. CD96 and CD200 have previously been identified as immunophenotypic markers of functional LSCs in adult AML(39, 40). In pediatric AML, high CD200 protein expression has been associated with poor clinical outcomes(56).

Patients who failed to achieve complete remission exhibited elevated CD200 levels, which were also linked to minimal residual disease. CD200 has been separately implicated in immune evasion in both adult(57) and pediatric AML(58). These shared immunophenotypic marker gene expressions further demonstrated the molecular overlap between scaLSC-SR cells and classical LSC phenotypes. Using SCA, we showed key features of similarity in LSC self-renewal programs between adult and pediatric and human AML. This highlighted conserved molecular mechanism that would lead to therapy resistance and disease relapse across different age groups in AML.

The mutational landscape is a major determinant of clinical and biological behavior in AML. However, the role of genetics in LSC behavior has not been well-studied. Expression of an LSC transcriptional signature has been shown to be prognostic in AML(6, 59), and is enriched in poor-risk genetic subtypes of AML(60, 61). Accordingly, we used deconvolution analysis and found that *TP53Mut* and *NRASMut* AMLs harbor significantly higher scaLSC-SR compositions compared to wildtype AMLs in two independent datasets. This finding is consistent with prior studies showing that both *TP53* and *NRAS* mutations enhance self-renewal capacity in AML (12, 13, 41, 42). Our group previously demonstrated, using a genetically engineered mouse model, that NRAS^G12V^ enforces a leukemia self-renewal gene expression program and is essential for leukemia self-renewal, independent of its effects on proliferation and survival(12, 41). Similarly, *TP53* mutations have been reported to confer gain-of-function effects that enhance LSC self-renewal capacity in mouse models(42). Mouse HSPCs engineered with the *Trp53^R172H^*mutation displayed unique transcriptional changes and enhanced self-renewal potential *in vivo*.

Finally, we demonstrated that the LSC-SR28 gene signature, derived from scRNA-seq data of human AML patients, is prognostic across four independent datasets. In addition, our LSC-SR28 signature scores were significantly higher in functionally validated LSC fractions compared to non LSC fractions, demonstrating that the LSC-SR28 score captured LSC stemness programs. Therefore, we showed that SCA reveals precise, clinically meaningful transcriptional behaviors and LSC-SR28 gene signature captures stemness program from AML patients.

## Acknowledgments

1. Z. Sachs was supported by the American Society of Hematology Scholar Award, American Cancer Society, Frederick A. DeLuca Foundation, Mentored Research Scholar Grant (MRSG-16-195-01-DDC); the Lois and Richard King Assistant Professorship in Medicine at the University of Minnesota, the Clinical and Translational Science Institute at the University of Minnesota KL2 Career Development Award and K to R01 Award (NIH/NCATS ULI RR033183 & KL2 RR0333182); the American Cancer Society Institutional Research Grant at the University of Minnesota (124166-IRG-58-001-55-IRG12); the Masonic Cancer Center at the University of Minnesota Translational Working Group Award and Genetic Mechanisms of Cancer Award; the University of Minnesota Department of Medicine Women’s Early Research Career Award; the division of Hematology, Oncology, and Transplantation, Department of Medicine; and the University of Minnesota Foundation donors. We extend our appreciation to the Minnesota Supercomputing Institute at the University of Minnesota.

## Author Contributions

YL conceptualized algorithm, designed the study, curated data, analyzed data, visualized data, created R software, and wrote the manuscript. WW conceptualized algorithm, designed the study, supervised data analysis, reviewed and edited manuscript. TKS provided resources. KEO provided resources, reviewed and edited manuscript. CLM designed the study concept, conceptualized algorithm, supervised the project, and wrote the manuscript. ZS designed the study concept, acquired funding, supervised the project, and wrote the manuscript.

Conflict-of-interest disclosure: The authors declare no competing financial interests.

Correspondence: Zohar Sachs, Mayo Mail Code 806, 420 Delaware St SE, Minneapolis, MN55455.

---

**Supplemental Figure S1.**
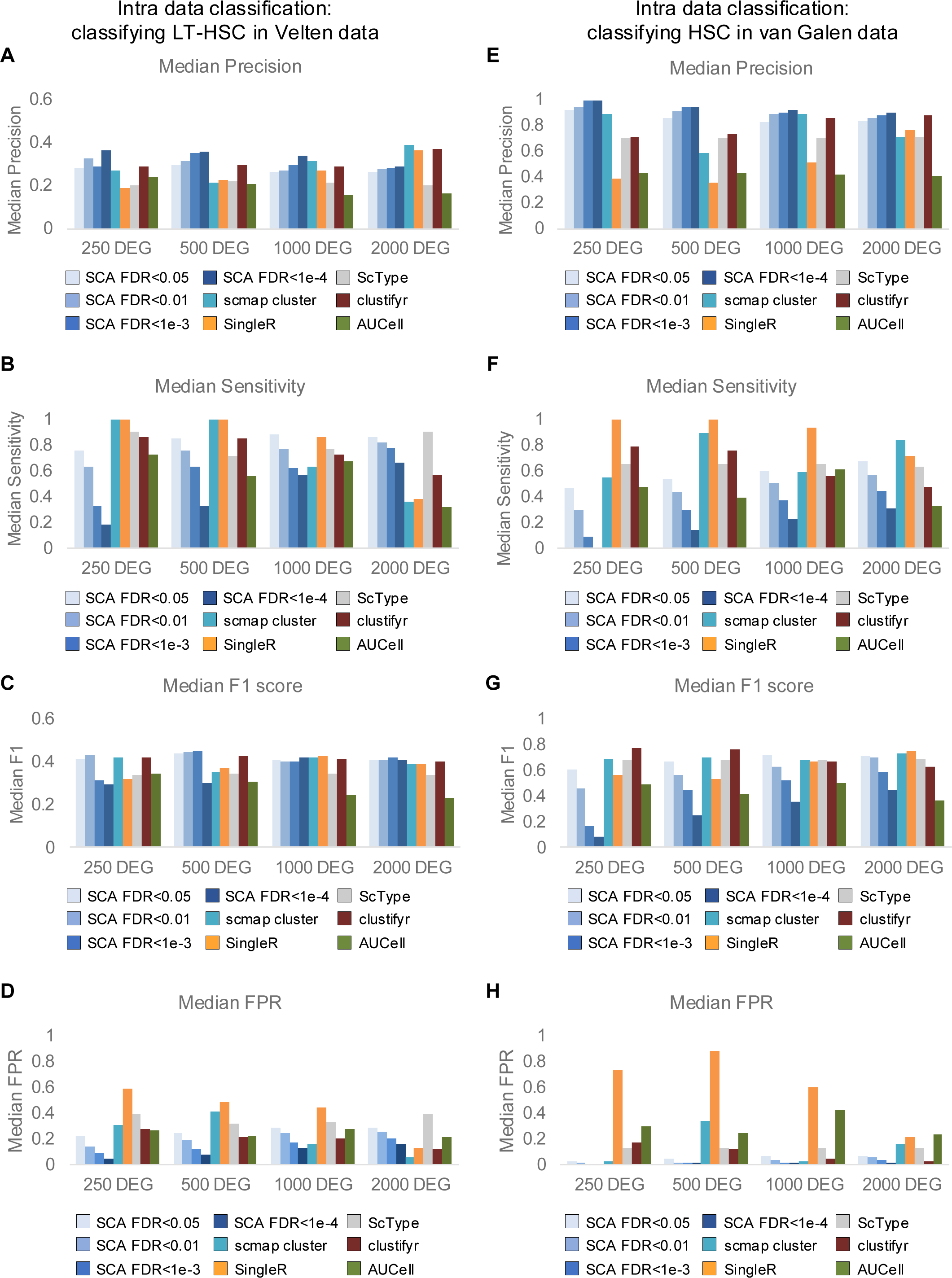

**Supplemental Figure S2.**
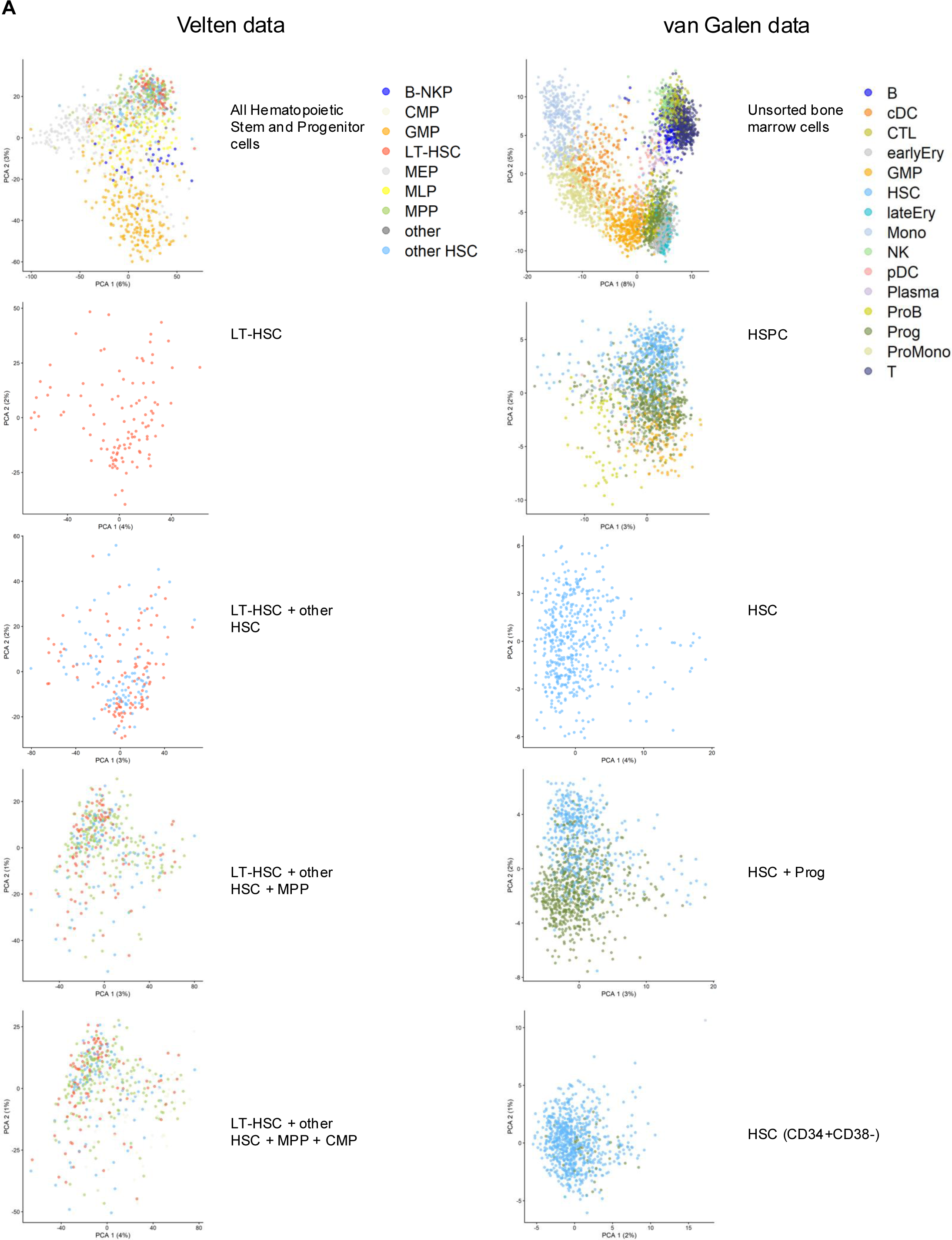
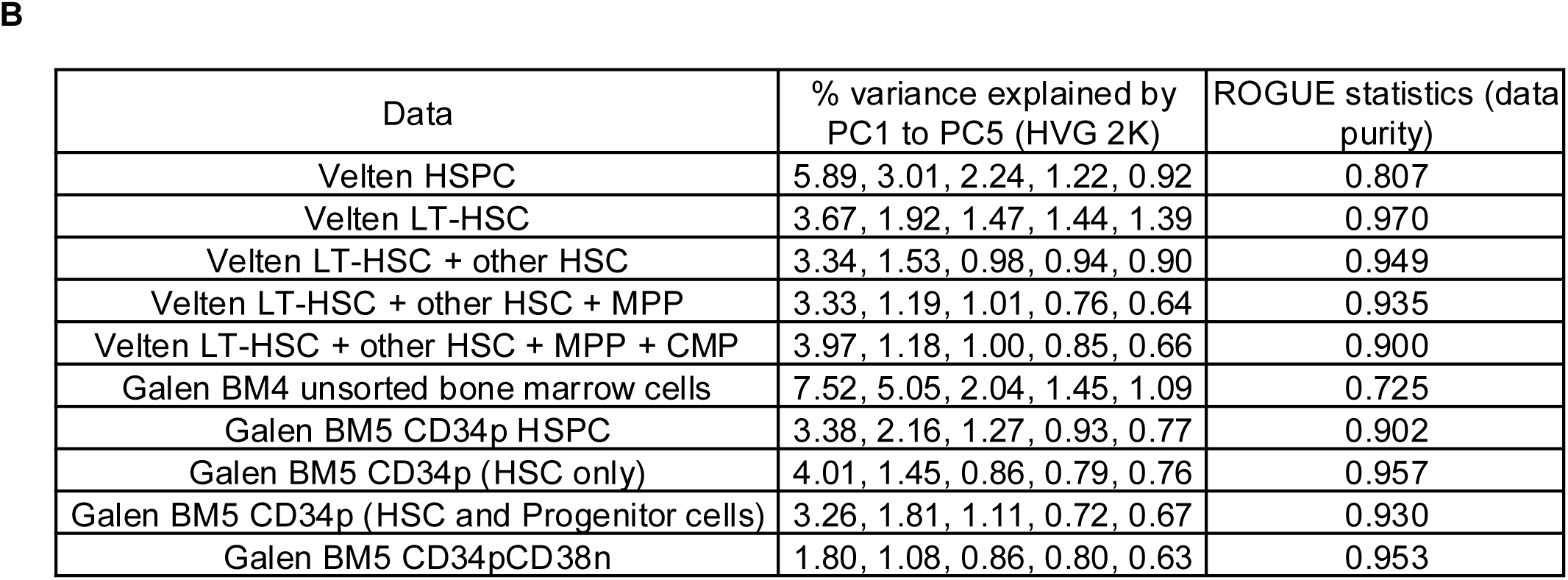

**Supplemental Figure S3.**
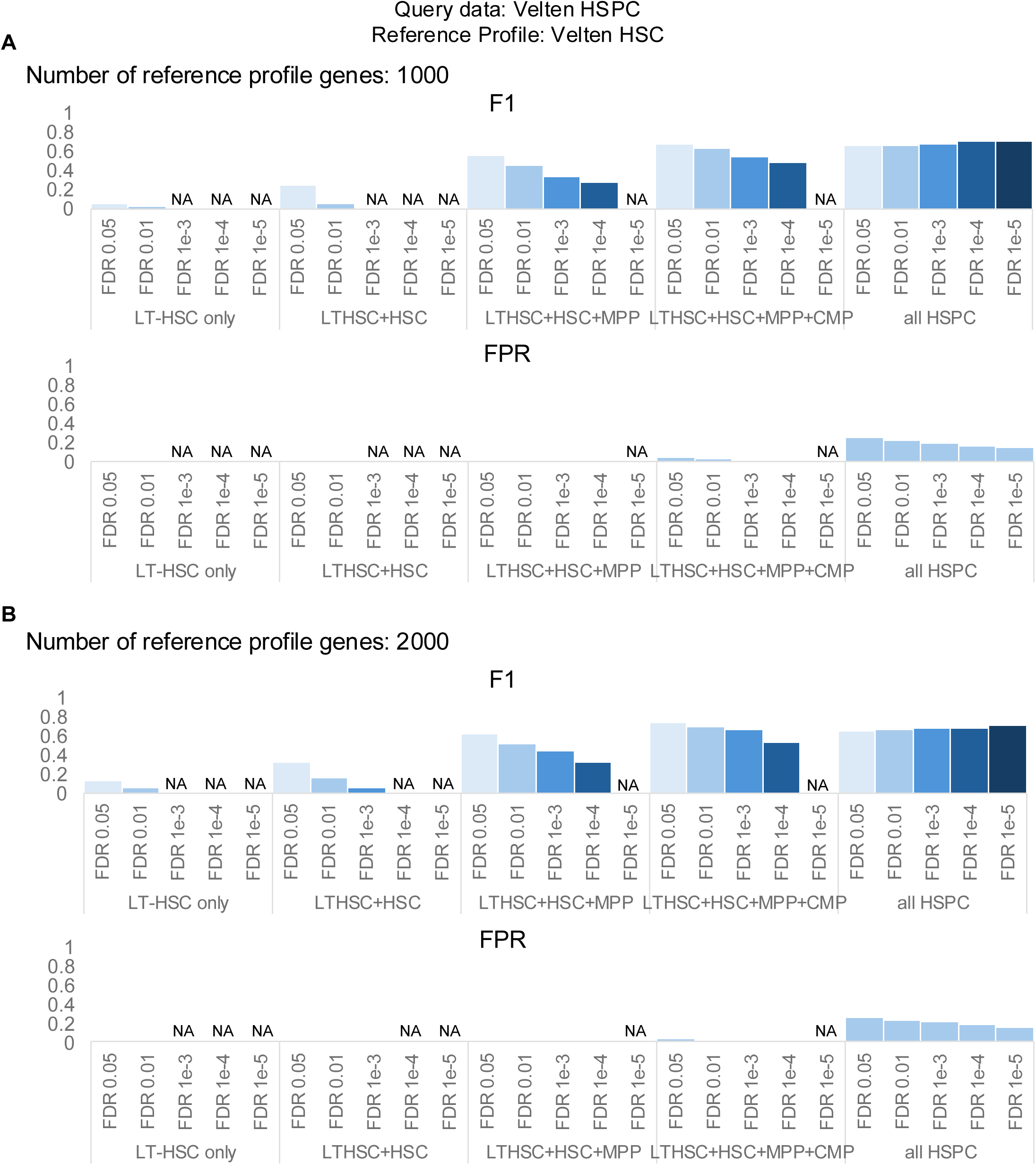

**Supplemental Figure S4.**
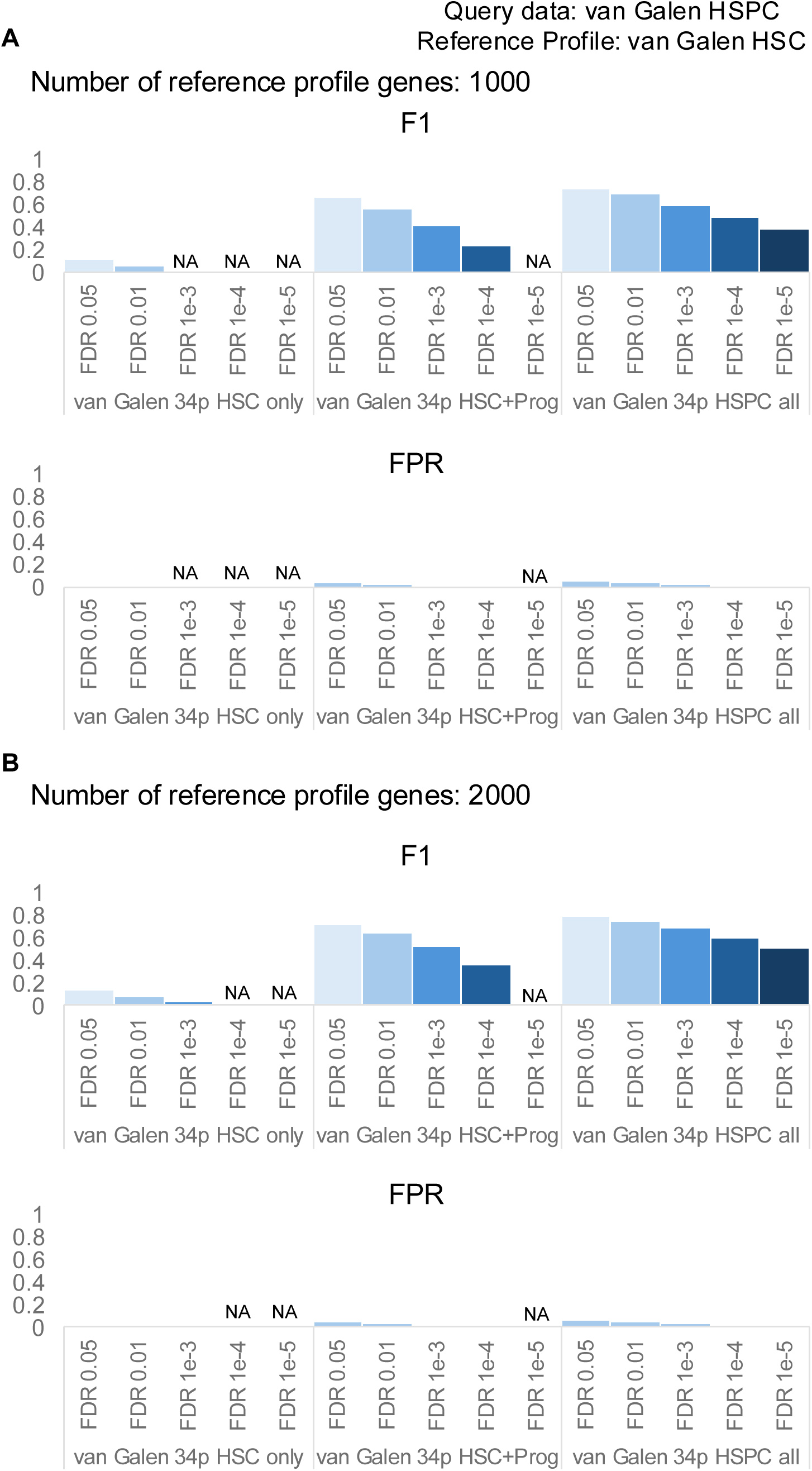

**Supplemental Figure S5.**
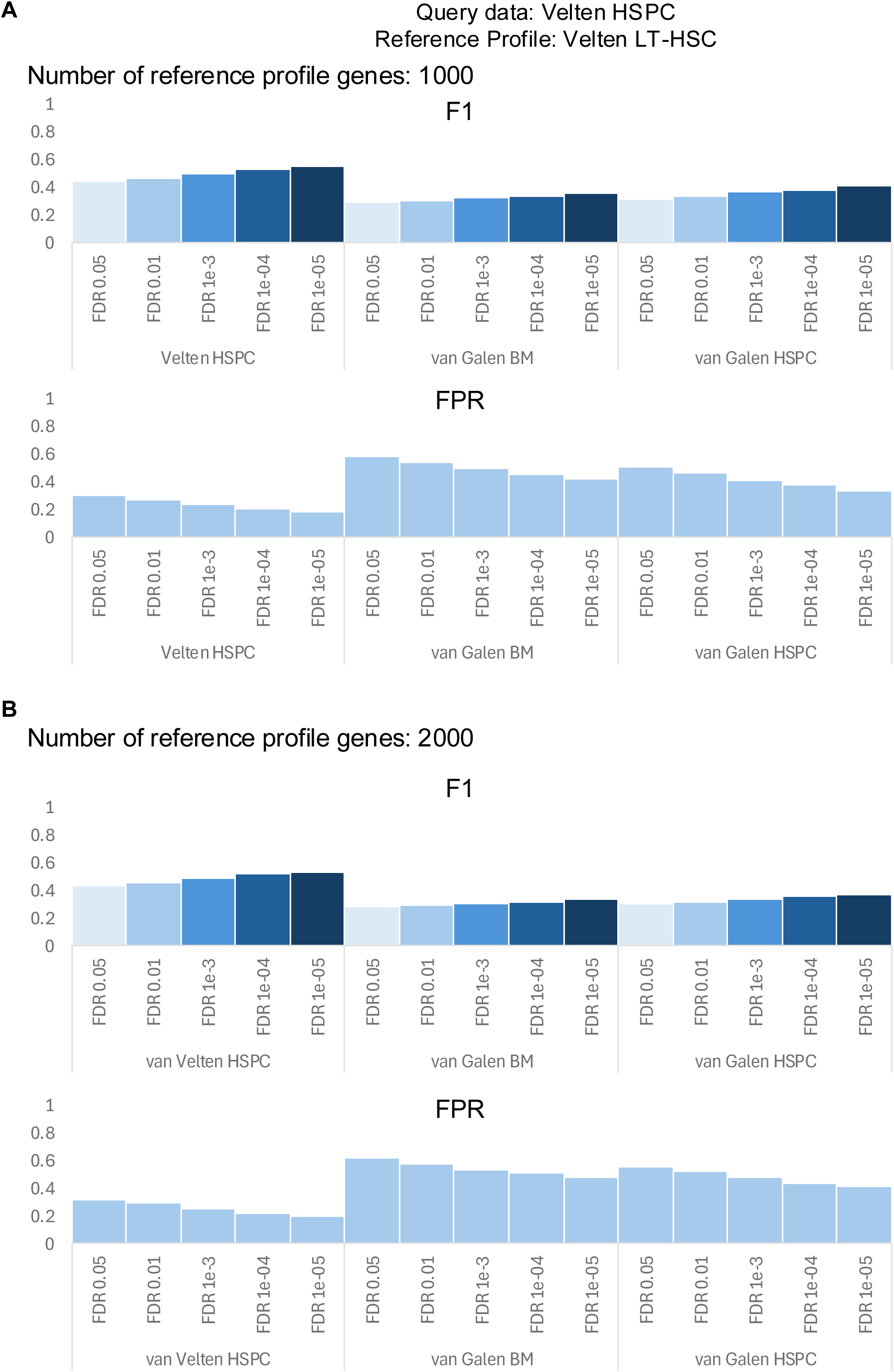

**Supplemental Figure S6.**
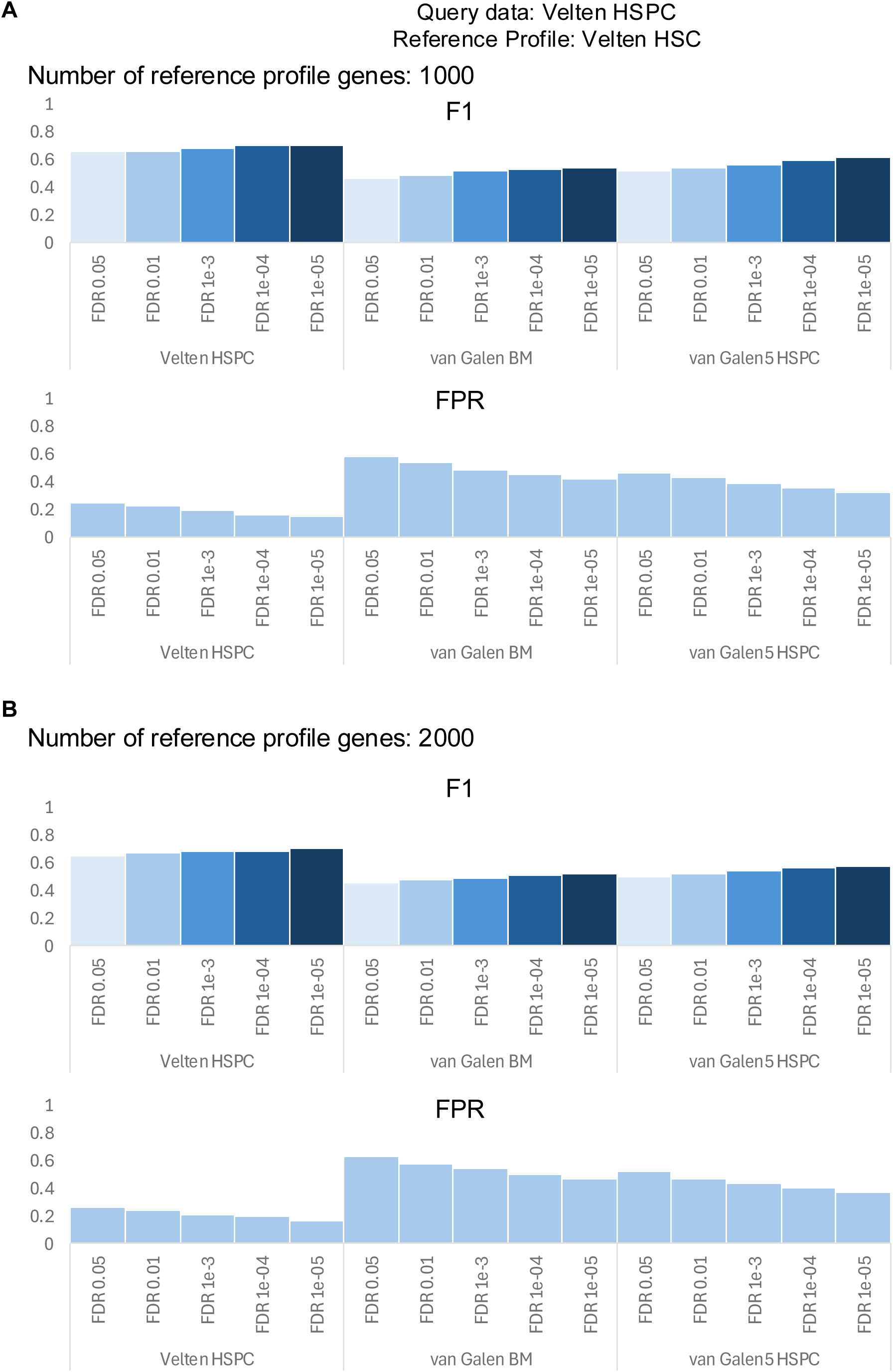

**Supplemental Figure S7.**
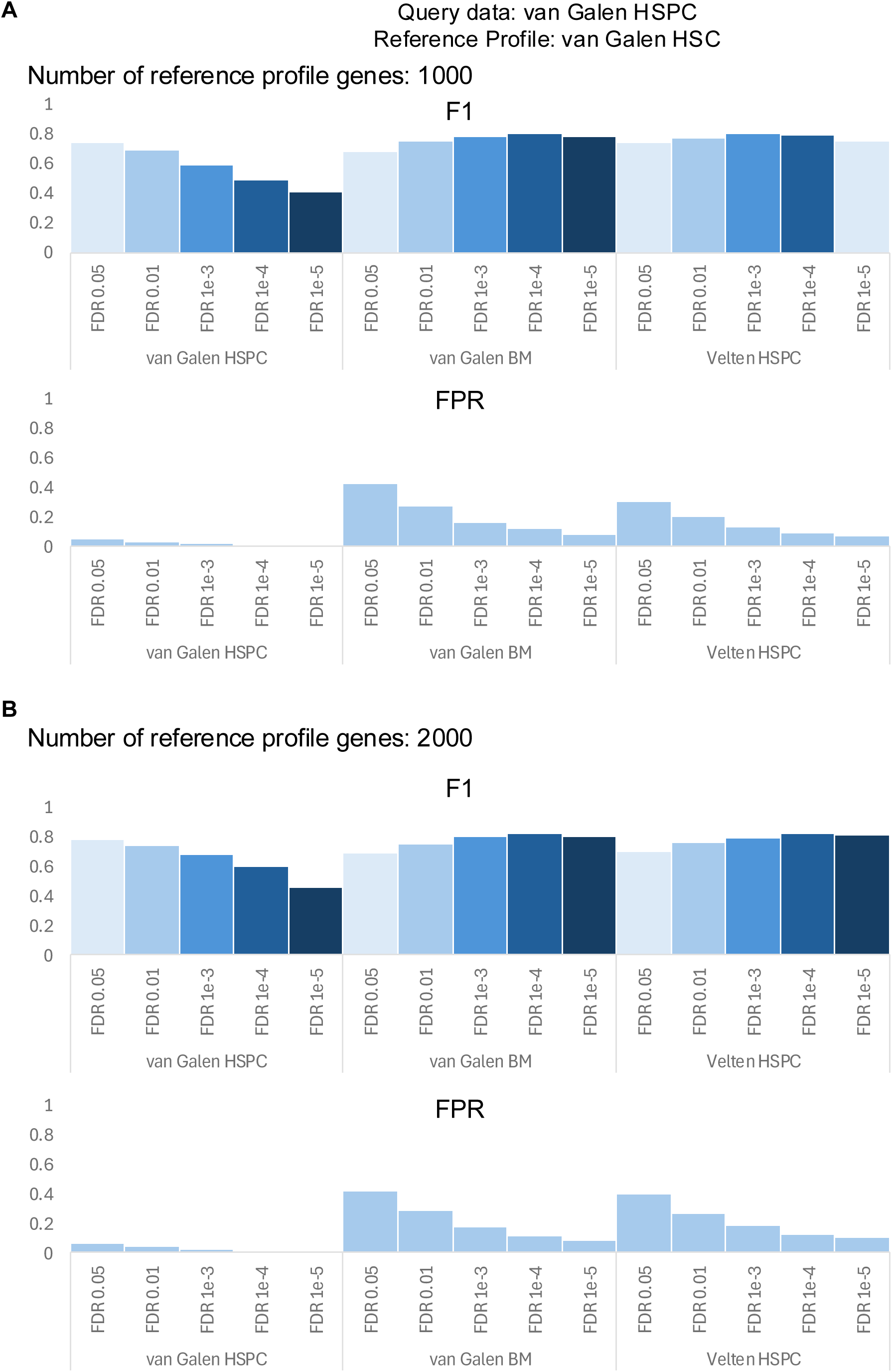

**Supplemental Figure S8.**
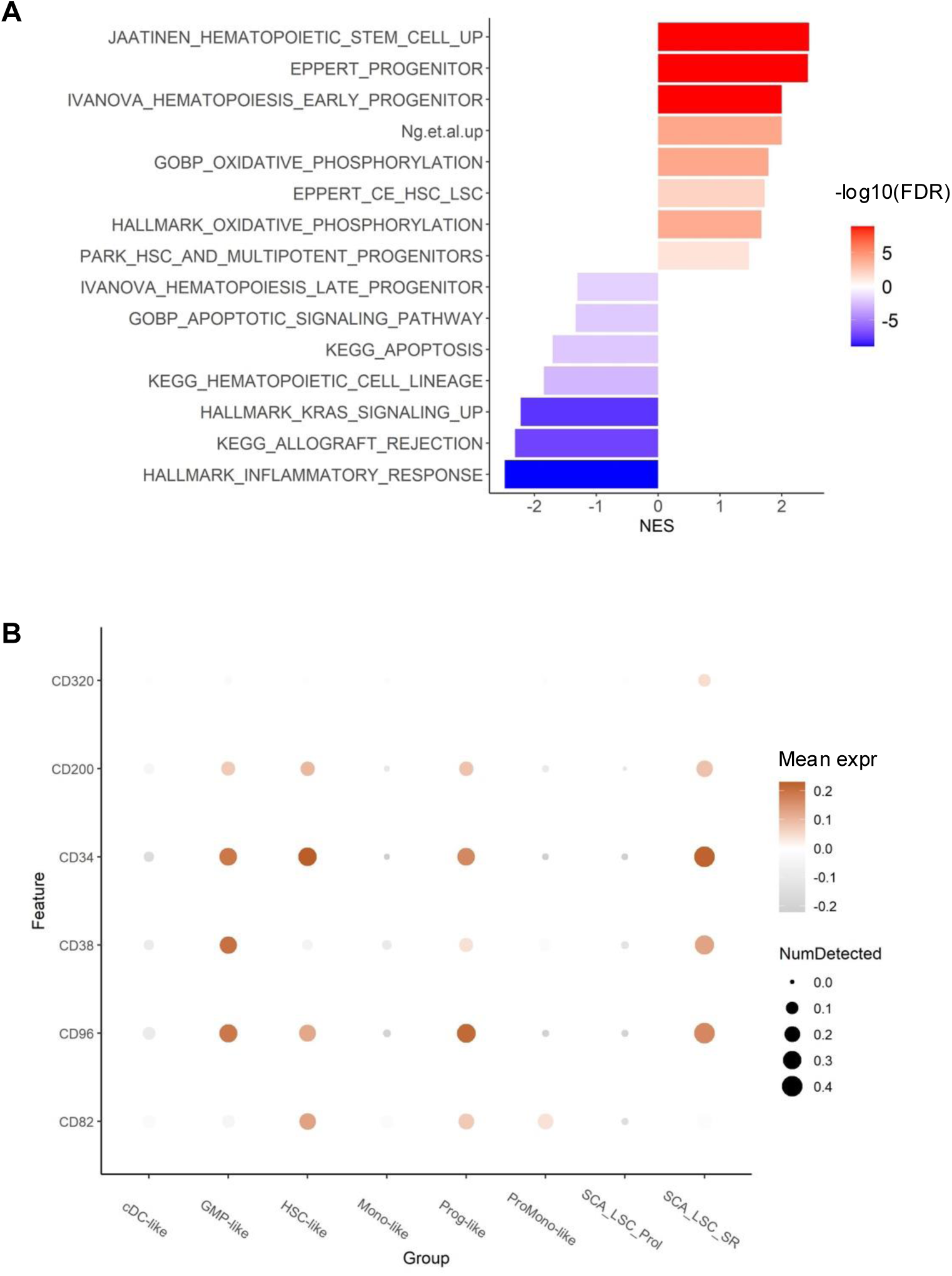

**Supplemental Figure S9.**
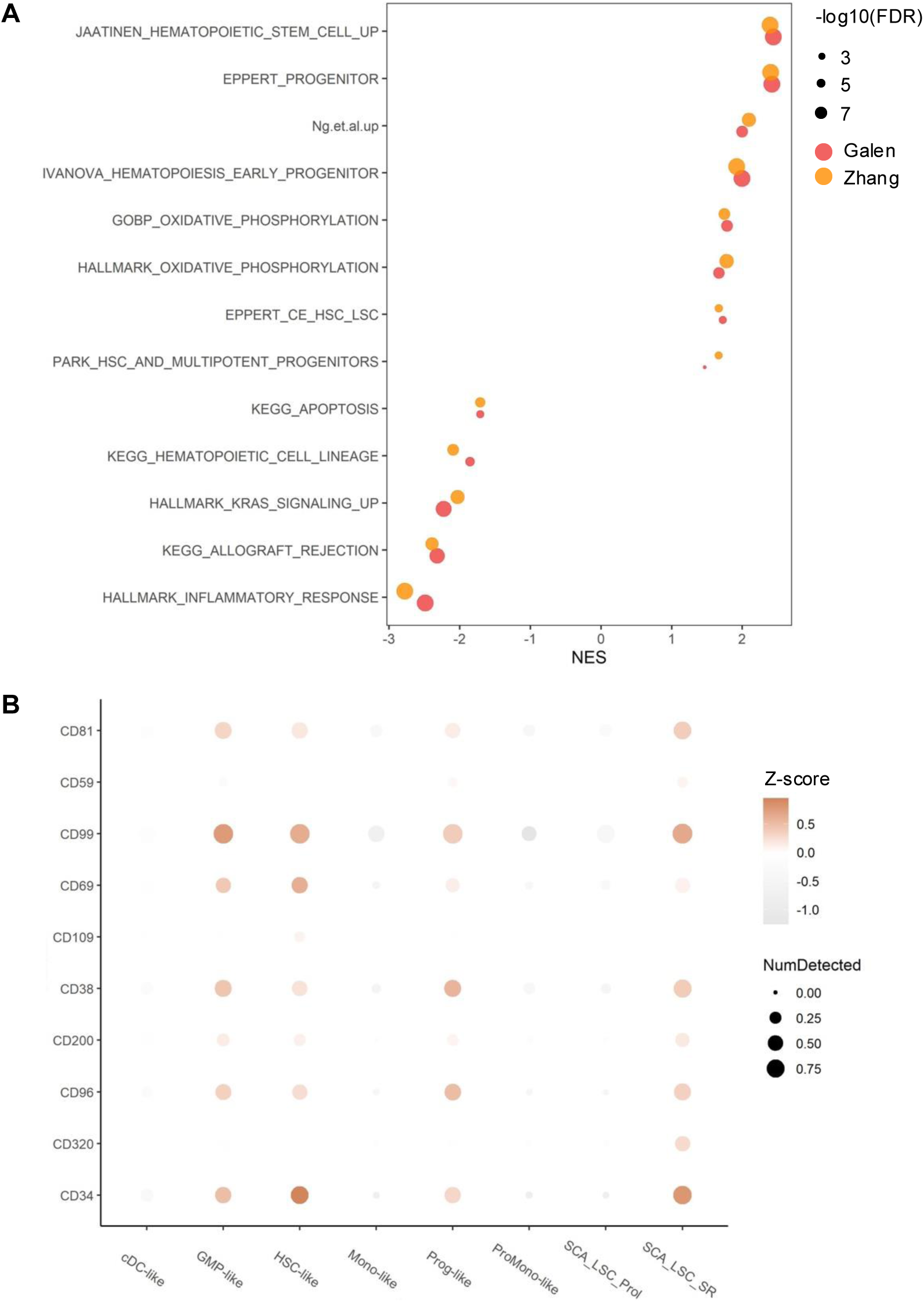

**Supplemental Figure S10.**
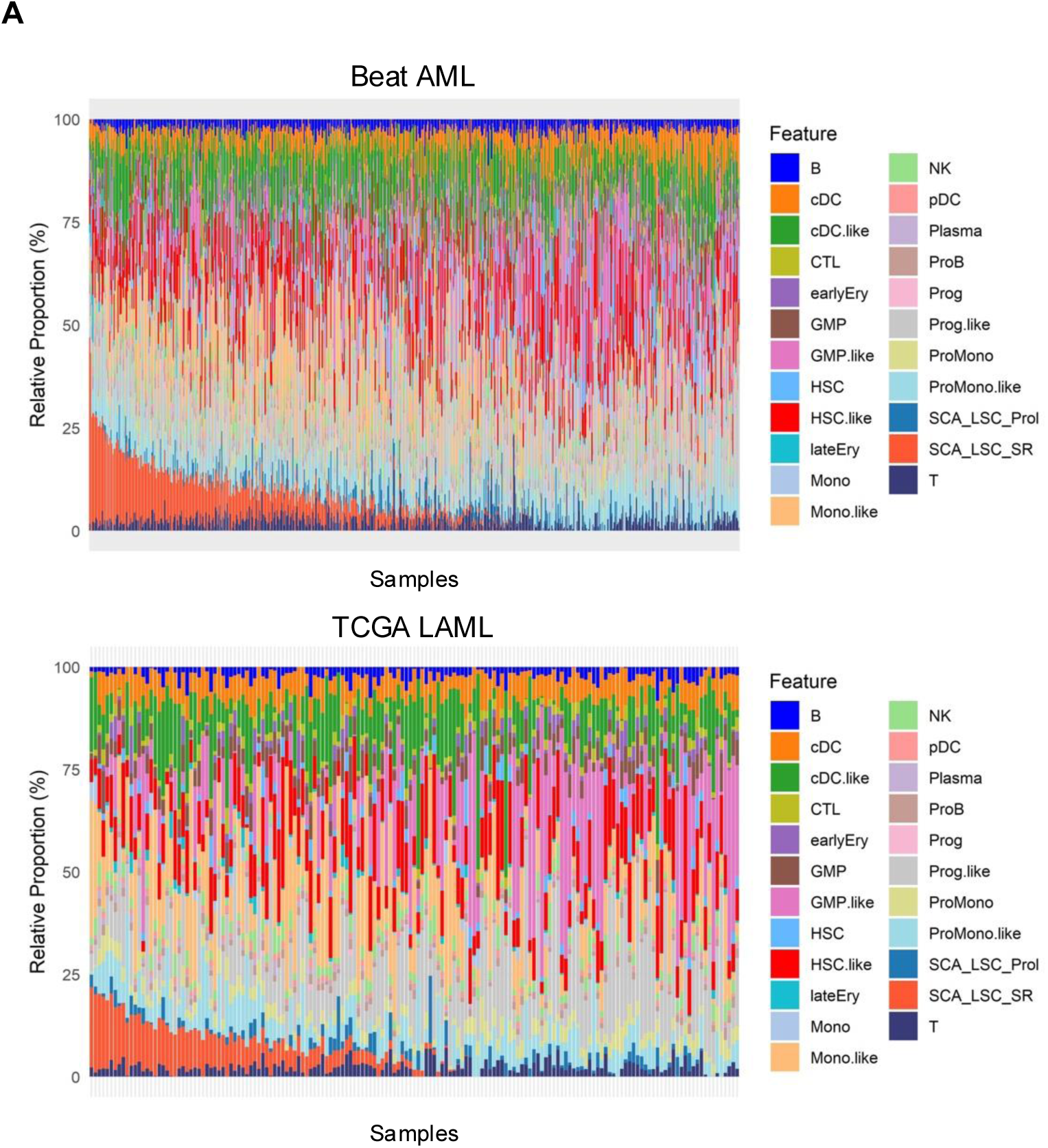
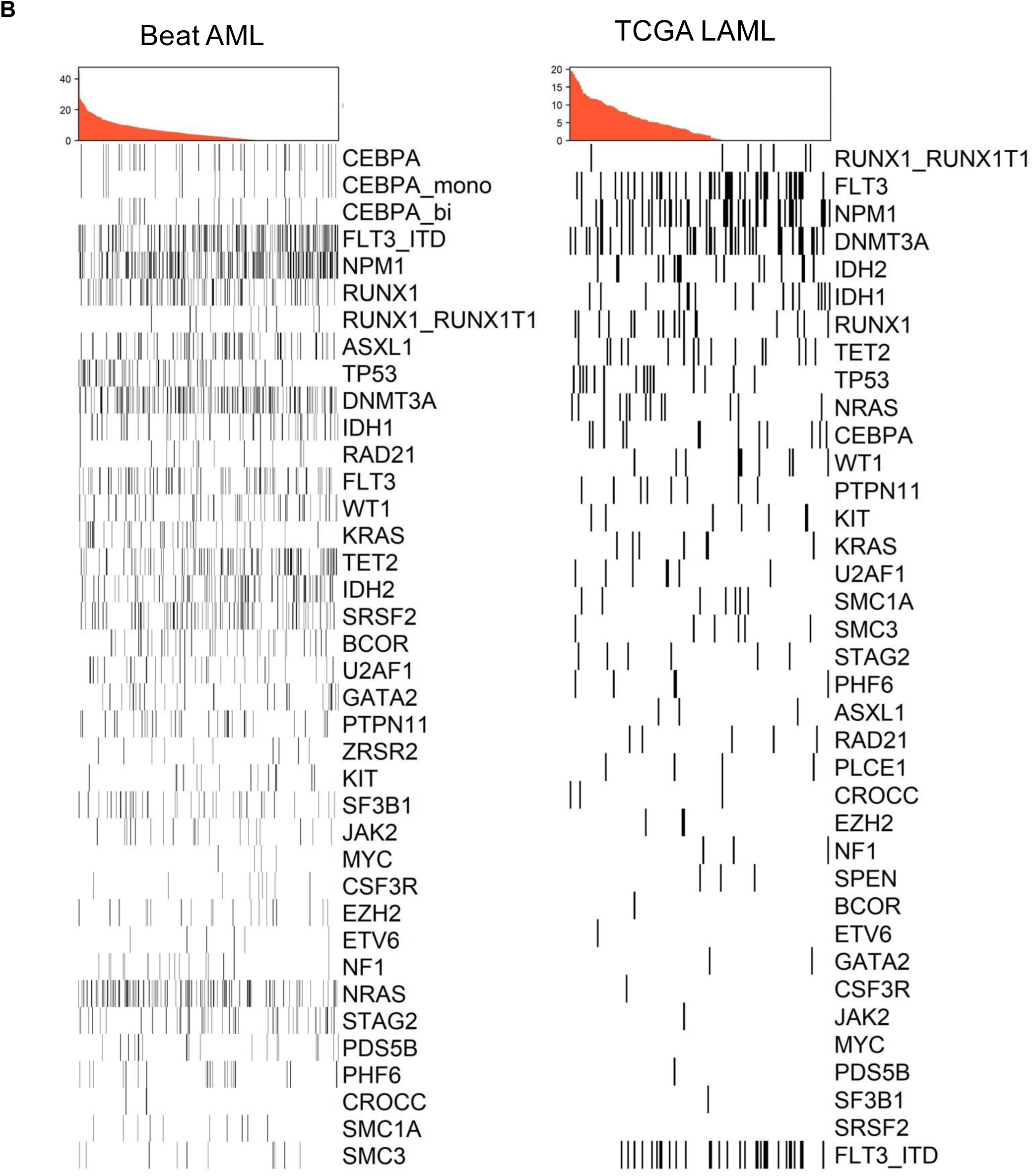

**Supplemental Figure S11.**
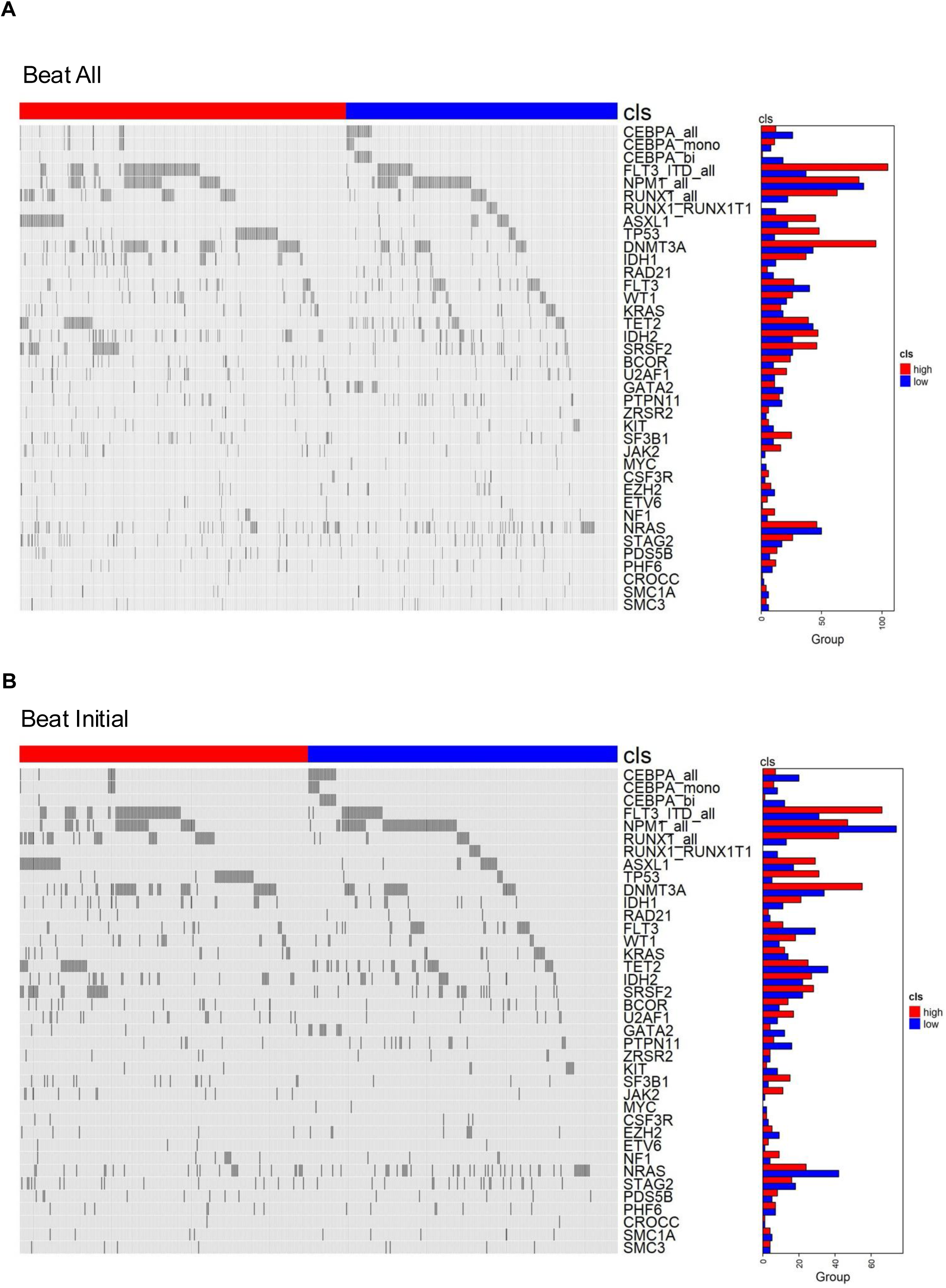
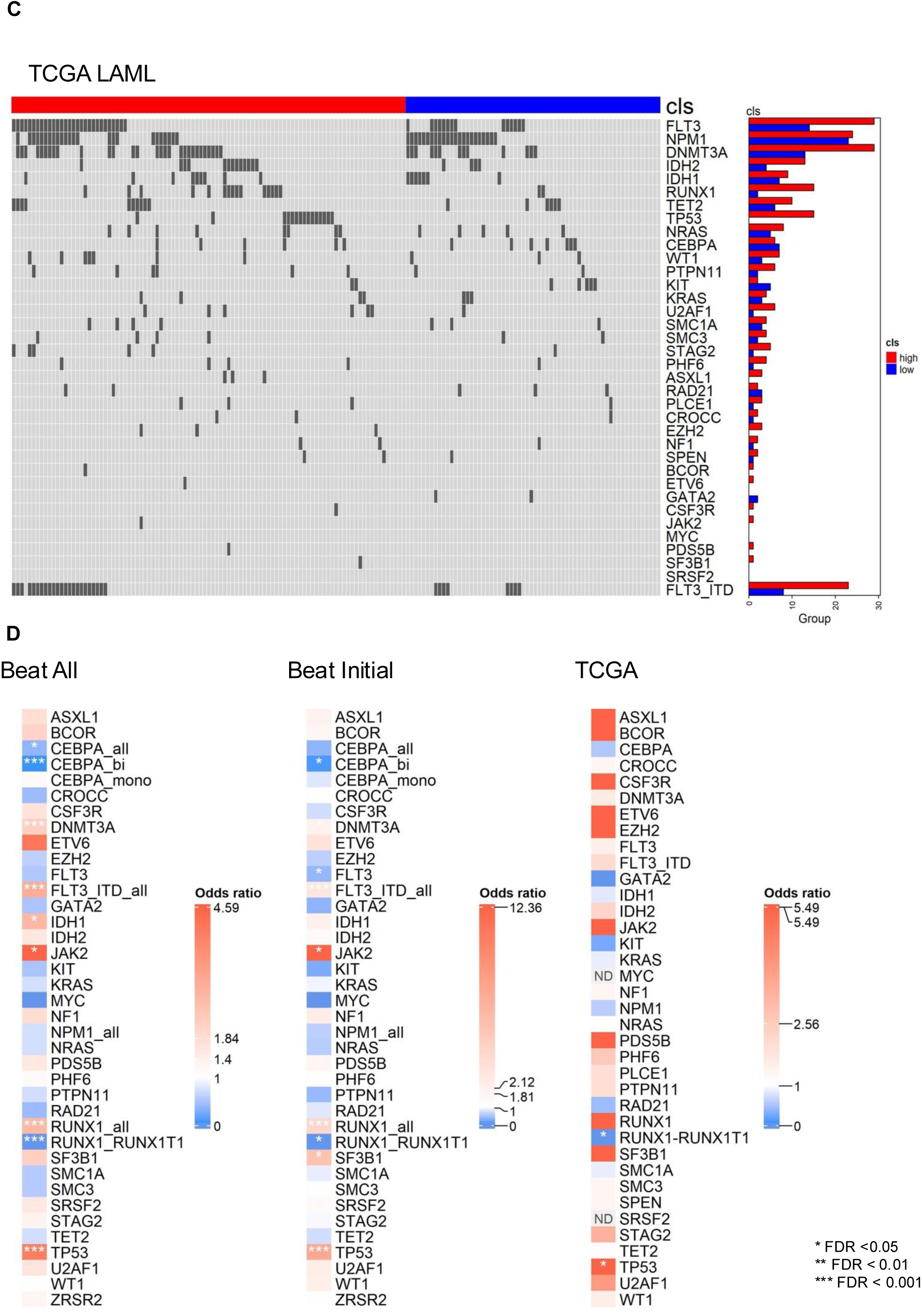

## Notes

### Competing Interest Statement

The authors have declared no competing interest.

## References

1. Metzeler KH, Herold T, Rothenberg-Thurley M, Amler S, Sauerland MC, Gorlich D, et al. Spectrum and prognostic relevance of driver gene mutations in acute myeloid leukemia. Blood. 2016;128(5):686–98.

2. Kubal T, Lancet JE. The thorny issue of relapsed acute myeloid leukemia. Current Opinion in Hematology. 2013;20(2):100–6.

3. Kreso A, Dick JE. Evolution of the cancer stem cell model. Cell Stem Cell. 2014;14(3):275–91.

4. Thomas D, Majeti R. Biology and relevance of human acute myeloid leukemia stem cells. Blood. 2017;129(12):1577–85.

5. Pollyea DA, Jordan CT. Therapeutic targeting of acute myeloid leukemia stem cells. Blood. 2017;129(12):1627–35.

6. Eppert K, Takenaka K, Lechman ER, Waldron L, Nilsson B, van Galen P, et al. Stem cell gene expression programs influence clinical outcome in human leukemia. Nat Med. 2011;17(9):1086–93.

7. Gentles AJ, Plevritis SK, Majeti R, Alizadeh AA. Association of a leukemic stem cell gene expression signature with clinical outcomes in acute myeloid leukemia. JAMA. 2010;304(24):2706–15.

8. Ng SW, Mitchell A, Kennedy JA, Chen WC, McLeod J, Ibrahimova N, et al. A 17-gene stemness score for rapid determination of risk in acute leukaemia. Nature. 2016;540(7633):433–7.

9. Zeng AGX, Bansal S, Jin L, Mitchell A, Chen WC, Abbas HA, et al. A cellular hierarchy framework for understanding heterogeneity and predicting drug response in acute myeloid leukemia. Nat Med. 2022;28(6):1212–23.

10. van Galen P, Hovestadt V, Wadsworth Ii MH, Hughes TK, Griffin GK, Battaglia S, et al. Single-Cell RNA-Seq Reveals AML Hierarchies Relevant to Disease Progression and Immunity. Cell. 2019;176(6):1265–81.e24.

11. Naldini MM, Casirati G, Barcella M, Rancoita PMV, Cosentino A, Caserta C, et al. Longitudinal single-cell profiling of chemotherapy response in acute myeloid leukemia. Nat Commun. 2023;14(1):1285.

12. Sachs K, Sarver AL, Noble-Orcutt KE, LaRue RS, Antony ML, Chang D, et al. Single-Cell Gene Expression Analyses Reveal Distinct Self-Renewing and Proliferating Subsets in the Leukemia Stem Cell Compartment in Acute Myeloid Leukemia. Cancer Res. 2020;80(3):458–70.

13. Sachs Z, LaRue RS, Nguyen HT, Sachs K, Noble KE, Mohd Hassan NA, et al. NRASG12V oncogene facilitates self-renewal in a murine model of acute myelogenous leukemia. Blood. 2014;124(22):3274–83.

14. Li Q, Bohin N, Wen T, Ng V, Magee J, Chen SC, et al. Oncogenic Nras has bimodal effects on stem cells that sustainably increase competitiveness. Nature. 2013;504(7478):143–7.

15. Alquicira-Hernandez J, Sathe A, Ji HP, Nguyen Q, Powell JE. scPred: accurate supervised method for cell-type classification from single-cell RNA-seq data. Genome Biology. 2019;20.

16. Aran D, Looney AP, Liu L, Wu E, Fong V, Hsu A, et al. Reference-based analysis of lung single-cell sequencing reveals a transitional profibrotic macrophage. Nature Immunology. 2019;20(2):163–72.

17. Kiselev VY, Yiu A, Hemberg M. scmap: projection of single-cell RNA-seq data across data sets. Nature Methods. 2018;15(5):359–62.

18. Pliner HA, Shendure J, Trapnell C. Supervised classification enables rapid annotation of cell atlases. Nature methods. 2019;16(10):983–6.

19. Tan Y, Cahan P. SingleCellNet: a computational tool to classify single cell RNA-Seq data across platforms and across species. Cell systems. 2019;9(2):207–13.e2.

20. Abdelaal T, Michielsen L, Cats D, Hoogduin D, Mei H, Reinders MJT, et al. A comparison of automatic cell identification methods for single-cell RNA sequencing data. Genome Biology. 2019;20(1):194.

21. Pasquini G, Rojo Arias JE, Schafer P, Busskamp V. Automated methods for cell type annotation on scRNA-seq data. Comput Struct Biotechnol J. 2021;19:961–9.

22. Velten L, Haas SF, Raffel S, Blaszkiewicz S, Islam S, Hennig BP, et al. Human haematopoietic stem cell lineage commitment is a continuous process. Nature cell biology. 2017;19(4):271–81.

23. Rodriguez-Fraticelli AE, Weinreb C, Wang S-W, Migueles RP, Jankovic M, Usart M, et al. Single-cell lineage tracing unveils a role for TCF15 in haematopoiesis. Nature. 2020;583(7817):585–9.

24. Zhang Y, Jiang S, He F, Tian Y, Hu H, Gao L, et al. Single-cell transcriptomics reveals multiple chemoresistant properties in leukemic stem and progenitor cells in pediatric AML. Genome Biology. 2023;24(1):199.

25. Bottomly D, Long N, Schultz AR, Kurtz SE, Tognon CE, Johnson K, et al. Integrative analysis of drug response and clinical outcome in acute myeloid leukemia. Cancer Cell. 2022;40(8):850–64 e9.

26. Cancer Genome Atlas Research N, Ley TJ, Miller C, Ding L, Raphael BJ, Mungall AJ, et al. Genomic and epigenomic landscapes of adult de novo acute myeloid leukemia. N Engl J Med. 2013;368(22):2059–74.

27. Rahman M, Jackson LK, Johnson WE, Li DY, Bild AH, Piccolo SR. Alternative preprocessing of RNA-Sequencing data in The Cancer Genome Atlas leads to improved analysis results. Bioinformatics. 2015;31(22):3666–72.

28. Tyner JW, Tognon CE, Bottomly D, Wilmot B, Kurtz SE, Savage SL, et al. Functional genomic landscape of acute myeloid leukaemia. Nature. 2018;562(7728):526–31.

29. Storey JD, Tibshirani R. Statistical significance for genomewide studies. Proc Natl Acad Sci U S A. 2003;100(16):9440–5.

30. Liu B, Li C, Li Z, Wang D, Ren X, Zhang Z. An entropy-based metric for assessing the purity of single cell populations. Nat Commun. 2020;11(1):3155.

31. McCarthy DJ, Chen Y, Smyth GK. Differential expression analysis of multifactor RNA-Seq experiments with respect to biological variation. Nucleic Acids Res. 2012;40(10):4288–97.

32. Subramanian A, Tamayo P, Mootha VK, Mukherjee S, Ebert BL, Gillette MA, et al. Gene set enrichment analysis: A knowledge-based approach for interpreting genome-wide expression profiles. Proceedings of the National Academy of Sciences. 2005;102(43):15545–50.

33. Newman AM, Steen CB, Liu CL, Gentles AJ, Chaudhuri AA, Scherer F, et al. Determining cell type abundance and expression from bulk tissues with digital cytometry. Nat Biotechnol. 2019;37(7):773–82.

34. Herold T, Jurinovic V, Batcha AMN, Bamopoulos SA, Rothenberg-Thurley M, Ksienzyk B, et al. A 29-gene and cytogenetic score for the prediction of resistance to induction treatment in acute myeloid leukemia. Haematologica. 2018;103(3):456–65.

35. Valk PJ, Verhaak RG, Beijen MA, Erpelinck CA, Barjesteh van Waalwijk van Doorn-Khosrovani S, Boer JM, et al. Prognostically useful gene-expression profiles in acute myeloid leukemia. N Engl J Med. 2004;350(16):1617–28.

36. Nader K, Tasci M, Ianevski A, Erickson A, Verschuren EW, Aittokallio T, et al. ScType enables fast and accurate cell type identification from spatial transcriptomics data. Bioinformatics. 2024;40(7).

37. Fu R, Gillen AE, Sheridan RM, Tian C, Daya M, Hao Y, et al. clustifyr: an R package for automated single-cell RNA sequencing cluster classification. F1000Res. 2020;9:223.

38. Aibar S, Gonzalez-Blas CB, Moerman T, Huynh-Thu VA, Imrichova H, Hulselmans G, et al. SCENIC: single-cell regulatory network inference and clustering. Nat Methods. 2017;14(11):1083–6.

39. Ho JM, Dobson SM, Voisin V, McLeod J, Kennedy JA, Mitchell A, et al. CD200 expression marks leukemia stem cells in human AML. Blood Adv. 2020;4(21):5402–13.

40. Hosen N, Park CY, Tatsumi N, Oji Y, Sugiyama H, Gramatzki M, et al. CD96 is a leukemic stem cell-specific marker in human acute myeloid leukemia. Proc Natl Acad Sci U S A. 2007;104(26):11008–13.

41. Kurata M, Antony ML, Noble-Orcutt KE, Rathe SK, Lee Y, Furuno H, et al. Proliferation and Self-Renewal Are Differentially Sensitive to NRASG12V Oncogene Levels in an Acute Myeloid Leukemia Cell Line. Mol Cancer Res. 2022;20(11):1646–58.

42. Loizou E, Banito A, Livshits G, Ho Y-J, Koche RP, Sánchez-Rivera FJ, et al. A Gain-of-Function p53-Mutant Oncogene Promotes Cell Fate Plasticity and Myeloid Leukemia through the Pluripotency Factor FOXH1. Cancer Discovery. 2019;9(7):962–79.

43. Dohner H, Wei AH, Appelbaum FR, Craddock C, DiNardo CD, Dombret H, et al. Diagnosis and management of AML in adults: 2022 recommendations from an international expert panel on behalf of the ELN. Blood. 2022;140(12):1345–77.

44. Haghverdi L, Lun ATL, Morgan MD, Marioni JC. Batch effects in single-cell RNA-sequencing data are corrected by matching mutual nearest neighbors. Nature Biotechnology. 2018;36(5):421–7.

45. Stuart T, Butler A, Hoffman P, Hafemeister C, Papalexi E, Mauck WM, et al. Comprehensive Integration of Single-Cell Data. Cell. 2019;177(7):1888–902.e21.

46. Stuart T, Satija R. Integrative single-cell analysis. Nature Reviews Genetics. 2019;20(5):257–72.

47. Ergen C, Xing G, Xu C, Kim M, Jayasuriya M, McGeever E, et al. Consensus prediction of cell type labels in single-cell data with popV. Nat Genet. 2024;56(12):2731–8.

48. Pei G, Yan F, Simon LM, Dai Y, Jia P, Zhao Z. deCS: A Tool for Systematic Cell Type Annotations of Single-cell RNA Sequencing Data among Human Tissues. Genomics, Proteomics & Bioinformatics. 2023;21(2):370–84.

49. Huang Q, Liu Y, Du Y, Garmire LX. Evaluation of Cell Type Annotation R Packages on Single-cell RNA-seq Data. Genomics, Proteomics & Bioinformatics. 2021;19(2):267–81.

50. Maan H, Zhang L, Yu C, Geuenich MJ, Campbell KR, Wang B. Characterizing the impacts of dataset imbalance on single-cell data integration. Nat Biotechnol. 2024;42(12):1899–908.

51. Ma W, Su K, Wu H. Evaluation of some aspects in supervised cell type identification for single-cell RNA-seq: classifier, feature selection, and reference construction. Genome Biol. 2021;22(1):264.

52. Bolouri H, Farrar JE, Triche T, Jr., Ries RE, Lim EL, Alonzo TA, et al. The molecular landscape of pediatric acute myeloid leukemia reveals recurrent structural alterations and age-specific mutational interactions. Nat Med. 2018;24(1):103–12.

53. Duployez N, Marceau-Renaut A, Villenet C, Petit A, Rousseau A, Ng SWK, et al. The stem cell-associated gene expression signature allows risk stratification in pediatric acute myeloid leukemia. Leukemia. 2019;33(2):348–57.

54. Elsayed AH, Rafiee R, Cao X, Raimondi S, Downing JR, Ribeiro R, et al. A six-gene leukemic stem cell score identifies high risk pediatric acute myeloid leukemia. Leukemia. 2020;34(3):735–45.

55. Stelmach P, Trumpp A. Leukemic stem cells and therapy resistance in acute myeloid leukemia. Haematologica. 2023;108(2):353–66.

56. Kandeel EZ, Madney Y, Eldin DN, Shafik NF. Overexpression of CD200 and CD123 is a major influential factor in the clinical course of pediatric acute myeloid leukemia. Exp Mol Pathol. 2021;118:104597.

57. Herbrich S, Baran N, Cai T, Weng C, Aitken MJL, Post SM, et al. Overexpression of CD200 is a Stem Cell-Specific Mechanism of Immune Evasion in AML. J Immunother Cancer. 2021;9(7).

58. Cieniewicz B, Uyeda MJ, Chen PP, Sayitoglu EC, Liu JM, Andolfi G, et al. Engineered type 1 regulatory T cells designed for clinical use kill primary pediatric acute myeloid leukemia cells. Haematologica. 2021;106(10):2588–97.

59. Ng SWK, Mitchell A, Kennedy JA, Chen WC, McLeod J, Ibrahimova N, et al. A 17-gene stemness score for rapid determination of risk in acute leukaemia. Nature. 2016;540(7633):433–7.

60. Bill M, Nicolet D, Kohlschmidt J, Walker CJ, Mrozek K, Eisfeld AK, et al. Mutations associated with a 17-gene leukemia stem cell score and the score’s prognostic relevance in the context of the European LeukemiaNet classification of acute myeloid leukemia. Haematologica. 2020;105(3):721–9.

61. Murphy T, Zhang B, Zhang T, King I, Capo-Chichi JM, Gupta V, et al. Real-world validation study of the LSC17 score for risk prediction in patients with newly diagnosed acute myeloid leukemia. Haematologica. 2025.

