## Supplemental Figure Legends for "Single cell Correlation Analysis (SCA): Identifying self-renewing subpopulation of human acute myeloid leukemia stem cells using single cell RNA sequencing analysis"

Supplemental Materials

Supplemental Figure S1. SCA has superior ability to generalize across datasets by optimizing false positive rates without compromising sensitivity and precision

Supplemental Figure S2. Cell population purity assessment using ROGUE statistic reflects expected biological diversity across hematopoietic background datasets

Supplemental Figure S3. Incorporating background data with higher cell type diversity improves the SCA performance in identifying HSCs in Velten HSPC dataset.

Supplemental Figure S4. Incorporating background data with higher cell type diversity improves the SCA performance in identifying HSCs in van Galen HSPC dataset.

Supplemental Figure S5. SCA can utilize background data from different samples, technology and capture methods in identifying LT-HSCs in Velten HSPC dataset.

Supplemental Figure S6. SCA can utilize background data from different samples, technology and capture methods in identifying HSCs in Velten HSPC dataset.

Supplemental Figure S7. SCA can utilize background data from different samples, technology and capture methods in identifying HSCs in van Galen HSPCs.

Supplemental Figure S8. Adult scaLSC-SR express self-renewal associated gene sets and cell surface marker encoding genes.

Supplemental Figure S9. Pediatric scaLSC-SR express self-renewal associated gene sets and cell surface marker encoding genes.

Supplemental Figure S10. Deconvolution analysis defined composition of scaLSC-SR in Beat AML and TCGA LAML.

Supplemental Figure S10. The composition of scaLSC-SR correlates with mutations in AML.

Supplemental Figure S11. scaLSC-SR gene signature defines distinct mutational subgroups in Beat AML and TCGA LAML patients.

Supplemental Figure Legends

Supplementary Figure 1. SCA has superior ability to generalize across datasets by optimizing false positive rates without compromising sensitivity and precision.

The ability of different scRNA-seq analysis methods, including SCA, to generalize was evaluated using a 5-fold cross-validation approach. Each dataset was randomly divided into five folds: one fold was used to generate the reference profile, while the remaining four were used for testing. This procedure was repeated five times, and the median performance metric was reported. To ensure consistency, the same folds were used across all tools in each test. Two independent human scRNA-seq datasets of normal human HSPCs were used for the query datasets: an experimentally validated Velten HSPC dataset and a second, independent dataset of van Galen HSPC data. The Velten data cell-type annotation is considered the gold-standard because these data are derived from index-sorted (i.e. flow cytometry-validated) cells. The van Galen data cell type annotation is based on clustering and marker gene. There are subtle differences in the annotation assignments across these two datasets. This analysis was conducted to classify LT-HSCs in the Velten dataset and HSCs in the van Galen dataset. To apply SCA and the other tools, Median precision (A, E), sensitivity (B, F), F1 score (C, G), and FPR (D, H) were measured as performance metrics. A-D LT-HSC classification in the Velten HSPC scRNA-seq dataset. In the Velten dataset, SCA with permissive FDR thresholds (FDR < 0.05 and 0.01) achieved the highest F1 scores across most reference profile sizes (C). When larger reference profiles (1,000 and 2,000 DEGs) were used, SCA consistently demonstrated comparable or superior F1 score across all FDR thresholds. In most cases, SCA with stringent FDR thresholds achieved the highest precision across reference profile sizes, except for the 2,000 DEG condition (A). Consistent with findings from inter-dataset and inter-species analyses, SCA using larger reference profiles generally exhibited lower sensitivity (B). However, with 2,000 DEGs, SCA maintained higher sensitivity than other methods tested. Furthermore, SCA with stringent FDR (FDR<10^-3^ and 10^-4^) achieved the lowest FPR across all reference profile sizes, except for the 2,000 DEGs condition (D). SCA with 2,000 DEGs showed the highest F1 scores across all thresholds. E-H HSC classification in the van Galen HSPC scRNA-seq dataset. HSCs in train fold was used to generate reference profile. The same train-test folds were used across all methods. Different numbers of differentially expressed genes were tested as reference profiles and the same reference profile was used across all tools tested. In the van Galen dataset, SCA with permissive FDR thresholds (FDR < 0.05 and 0.01) achieved F1 scores comparable to those of other methods (G). Across all tested reference profile sizes, SCA demonstrated the highest precision relative to other methods (E), although SCA generally exhibited lower sensitivity (F). SCA achieved the lowest FPR under permissive FDR thresholds (FDR < 0.05 and 0.01), while still maintaining comparable F1 scores (E and G).

**Supplemental Figure S2. Cell population purity assessment using ROGUE statistic reflects expected biological diversity across hematopoietic background datasets.** **A** Principal component analysis (PCA) was performed on different background datasets, including Velten HSPCs, van Galen unsorted bone marrow (BM) cells, van Galen HSPCs, and van Galen HSCs. Van Galen unsorted BM cells and HSPCs/HSCs were obtained from two different donors. Velten and van Galen HSPC datasets were *in silico* sorted based on previously annotated cell types. PCA was conducted using 2,000 highly variable genes, and the percentages of variance explained by PC1 to PC5 were calculated. **B** Cell population purity for each background dataset was assessed using the ROGUE statistic. A ROGUE value near 1 indicates high cell population purity, while a value near 0 reflects high heterogeneity of the cell population.

Supplemental Figure S3. Incorporating background data with higher cell type diversity improves the SCA performance in identifying HSCs in Velten HSPC dataset. The impact of using different background datasets with varying levels of cell type diversity was evaluated. SCA was performed on Velten HSPCs as query data to identify HSCs using the query data’s HSCs as the reference profile with A 1000 genes and B 2000 genes. F1 scores and FPRs were calculated for HSC classification using different background datasets. Background datasets were created by sequentially adding cell populations from the query Velten data: LT-HSCs alone; LT-HSCs + other HSCs; LT-HSCs + other HSCs + MPPs; LT-HSCs + other HSCs + MPPs + CMPs; and all HSPCs. F1 scores and FPRs were measured at various FDR thresholds for each background dataset.

Supplemental Figure S4. Incorporating background data with higher cell type diversity improves the SCA performance in identifying HSCs in van Galen HSPC dataset.

The impact of using different background datasets with varying levels of cell type diversity was evaluated. SCA was performed on van Galen HSPCs as query data to identify HSCs using the query data’s HSCs as the reference profile with A 1000 genes and B 2000 genes. F1 scores and FPRs were calculated for HSC classification using different background datasets. Background datasets were created by sequentially adding cell populations from the query van Galen data: HSCs alone; HSCs + Progenitors; and all HSPCs. F1 scores and FPRs were measured at various FDR thresholds for each background dataset.

Supplemental Figure S5. SCA can utilize background data from different samples, technology and capture methods in identifying LT-HSCs in Velten HSPC dataset.

The impact of using background datasets derived from different samples, technologies, and capture methods was evaluated. SCA was performed on Velten HSPCs as query data to identify LT-HSCs using the query data’s LT-HSCs as the reference profile with A 1000 genes and B 2000 genes. F1 scores and FPRs were calculated for LT-HSC classification using different background datasets. The following background dataset were used: Velten HSPC dataset; van Galen unsorted bone marrow (BM) cells; and van Galen HSPCs.

Supplemental Figure S6. SCA can utilize background data from different samples, technology and capture methods in identifying HSCs in Velten HSPC dataset.

The impact of using background datasets derived from different samples, technologies, and capture methods was evaluated. SCA was performed on Velten HSPCs as query data to identify HSCs using the query data’s HSCs as the reference profile with A 1000 genes and B 2000 genes. F1 scores and false positive rates (FPR) were calculated for HSC classification using different background datasets. The following background dataset were used: Velten HSPC dataset; van Galen unsorted bone marrow (BM) cells; and van Galen HSPCs.

Supplemental Figure S7. SCA can utilize background data from different samples, technology and capture methods in identifying HSCs in van Galen HSPCs.

The impact of using background datasets derived from different samples, technologies, and capture methods was evaluated. SCA was performed on van Galen HSPCs as query data to identify HSCs using the query data’s HSCs as the reference profile with A 1000 genes and B 2000 genes. F1 scores and false positive rates (FPR) were calculated for HSC classification using different background datasets. The following background dataset were used: van Galen’s HSPC dataset itself; van Galen’s unsorted BM cells; and Velten’s HSPC dataset.

Supplemental Figure S8. Adult scaLSC-SR express self-renewal associated gene sets and cell surface marker encoding genes.

A GSEA was performed to compare scaLSC-SR cells with all the other AML cells. B Genes encoding cell surface markers that are differentially expressed between scaLSC-SR and other AML cells across all the samples with an FDR<0.05 are shown. Data are displayed as log2 transformed CPM expression values that were mean-centered (Z-score). Previously defined AML blast cell types (*-like* cells) and SCA-defined cell types were compared.

Figure 6. SCA identifies pediatric AML cells expressing the mouse LSC self-renewal and proliferation single cell gene expression profiles.

**A** tSNE visualization of 13 pediatric human AML samples’ scRNA-seq data (Zhang et al. Genome Biology 2022) is shown. Cell type annotation was transferred to the pediatric dataset using the AML scRNA-seq data from van Galen et al 2019 using SingleR (left). SCA-identified AML cells expressing the mouse LSC self-renewal (scaLSC-SR, red) and LSC proliferation (scaLSC-Pro, blue) single cell gene expression profiles with FDR< 1x10^-6^ (right). **B** Pie charts display cell type compositions in all samples in the dataset (left), within scaLSC-SR cells (middle), and within scaLSC-Prol cells (right). The percentages of each cell type are included in the graph legends. **C** Bar plots showing cell type compositions within each AML sample. Fractions of cell types are displayed for each sample (top left); along with fractions of scaLSC-SR and scaLSC-Pro cells in each sample (top right). The lower plots show cell type fractions within scaLSC-SR (bottom left) and within scaLSC-Pro (bottom right) across samples.

Supplemental Figure S9. Pediatric scaLSC-SR express self-renewal associated gene sets and cell surface marker encoding genes.

A GSEA was performed to compare scaLSC-SR cells with other AML cells in the scRNA-seq data and compared to the GSEA results from adult AML dataset. Gene sets displayed are those that are significantly enriched in both adult and pediatric scaLSC-SR cells, based on concordant normalized enrichment scores (NES) and FDR <0.05 in both datasets independently. B Genes encoding cell surface markers that are differentially expressed between scaLSC-SR and other AML cells across all the samples with an FDR<0.05 are shown. Data are displayed as log2-transformed CPM expression values that were mean-centered (Z-score). Previously defined AML blast cell types (*-like* cells) and SCA-defined cell types were compared.

Supplemental Figure S10. Deconvolution analysis defined composition of scaLSC-SR in Beat AML and TCGA LAML.

Deconvolution approach was applied to bulk RNAseq data from Beat AML and TCGA LAML to determine the cell-type composition of each sample in the two databases using the cell type annotation defined in van Galen 2019. A Samples (row) were sorted based on the abundance of scaLSC-SR compositions. B Mutational landscape of the most frequently mutated genes in the Beat AML (left) and TCGA LAML (right) datasets. Histograms in the upper panels show the percentages of scaLSC-SR identified in each sample.

Supplemental Figure S11. scaLSC-SR gene signature defines distinct mutational subgroups in Beat AML and TCGA LAML patients.

**A-C** Mutational landscape of the most frequently mutated genes of samples with high and low scaLSC-SR signature scores in **A** all Beat AML samples and **B** Beat AML diagnostic samples. Histograms on the right panels show the number of cases within each group **C** TCGA LAML (which includes only diagnostic samples). Histograms on the right panels show the number of cases within each group. **D** Enrichment analysis was conducted to compare odds ratio and p-value for enrichment or depletion of each mutation in high versus low LSC-Sr signature groups. Fisher’s exact test was used for enrichment analysis. Multiple hypothesis testing was corrected using Benjamini-Hochberg to calculate FDR. Asterisks denote mutations with statistically significant enrichment or depletion (FDR<0.05) in scaLSC-SR signature high group compared to low group.
